# 3D Microcapsules for Human Bone Marrow Derived Mesenchymal Stem Cell Biomanufacturing in a Vertical-Wheel Bioreactor

**DOI:** 10.1101/2023.02.16.528656

**Authors:** Matthew Teryek, Pankaj Jadhav, Raphaela Bento, Biju Parekkadan

## Abstract

Microencapsulation of human mesenchymal stromal cells (MSCs) via electrospraying has been well documented in tissue engineering and regenerative medicine. Herein, we report the use of microencapsulation, via electrospraying, for MSC expansion using a commercially available hydrogel that is durable, optimized to MSC culture, and enzymatically degradable for cell recovery. Critical parameters of the electrospraying encapsulation process such as seeding density, correlation of microcapsule output with hydrogel volume, and applied voltage were characterized to consistently fabricate cell-laden microcapsules of uniform size. Upon encapsulation, we then verified ∼ 10x expansion of encapsulated MSCs within a vertical-wheel bioreactor and the preservation of critical quality attributes such as immunophenotype and multipotency after expansion and cell recovery. Finally, we highlight the genetic manipulation of encapsulated MSCs as an example of incorporating bioactive agents in the capsule material to create new compositions of MSCs with altered phenotypes.

## Introduction

Human mesenchymal stromal cell (MSC) based therapies are of clinical interest due to their immunomodulatory properties through broad-spectrum release of trophic factors, multipotent differentiation capabilities, and low immunogenicity enabling an “off-the-shelf” allogeneic product [1-3]. The interventional therapeutic potential of these cell-based therapies has been investigated to alleviate an array of clinical indications that include hematopoietic failure [4-6], liver failure [7], multiple sclerosis [8], graft versus host disease [9, 10], and diabetes [11]. Alternative approaches using MSC therapeutics have gained traction, particularly in the *ex-vivo* genetic modification of MSCs [12] and MSC-derived exosomes [13-15], however both are in early phase development. Depending on the clinical indication, a single dose can range from 0.5×10^6^ to 5.0×10^6^ cells/kg of body weight [16]. When this dose is scaled for repeated administration per patient, the number of total patients per indication, and multiple jurisdictions of use, the cell mass required for commercial use approaches the order of 10^12^-10^13^ MSCs per year for a single indication as projected by Olsen *et al*. [17].

A major bottleneck in translating MSC therapies lies in manufacturing cell lots at a commercial scale to meet this clinical demand. Dosage requirements suggest that 2D cell culture is inadequate in addressing these demands without incurring tremendous costs, labor, and facility utilization [18]. Suspension-based culture is a feasible approach that utilizes automated stirred-tank systems to expand cells in 3D conditions while also being continuously monitored. Microcarriers, a well-researched scalable platform extensively used in the expansion of adherent MSCs, offer a larger surface area to volume ratio capable of supporting denser cultures once translated into suspension-based systems [19, 20]. Scaled-down pilot studies have shown that several microcarrier attributes such as size, porosity, chemically functionalized or extracellular coatings play a crucial role in cell expansion with implications in scaled-up commercial runs [21]. Notably these implications can dictate bioreactor agitation rates to not only keep microcarriers in suspension, but also for downstream processing. With too high an agitation rate having the potential to affect the desired target product profile, reduced cell health, or lead to particulate debris in final products as a regulated safety concern [22, 23].

Encapsulation of cells within 3D biocompatible polymer matrices has been investigated as an alternative to microcarrier based platforms. Such polymer matrices offer a stable and conducive platform for cell attachment and expansion within a porous material while allowing sufficient gas and nutrient exchange in suspension culture. Alginate, a natural biomaterial, has previously been reported to act as a medium for sustained drug delivery [24] and has frequently been associated with MSC microencapsulation [25-27]. Alginate however lacks adhesive moieties and is inherently unstable as chelating agents within culture media tend to displace the divalent crosslinker ionic interactions over time [28, 29]. In addition, biocompatible polymer-based encapsulation platforms come with their own set of challenges that include reports of hydrogel matrices leading to undesirable changes in cell functionality and premature differentiation of cells [30, 31].

In this study, we developed a microcapsule-based platform using VitroGel-MSC, a xeno-free polysaccharide-based hydrogel to expand MSCs in a vertical-wheel bioreactor following a fed-batch approach. Critical process parameters were evaluated and optimized to arrive at a multifold expansion of MSCs without compromising functionality and differentiating capabilities. Critical quality attributes (CQAs) were validated using multiple analytical techniques. Notably, we were also able to prove the functionality of our 3D biomanufacturing system by employing it as a platform to generate genetically manipulated MSCs via viral transduction. Herein, we demonstrate that this 3D cell expansion is feasible and serves as a proof-of-concept that can be considered for further scale-up and process development for MSC therapy biomanufacturing.

## Materials and Methods

### MSC planar culture for seed train

Human mesenchymal stromal cells (MSCs) were isolated from single donor bone marrow (Lonza, Walkersville, MD, USA) based upon their adherence to tissue-culture treated flasks in standard conditions. MSCs were cultured in minimal essential media-α (Thermo Fisher Scientific, Waltham, MA, USA) supplemented with 2.5 ng/mL rec. human FGF-2 (Waisman Biomanufacturing, Madison, WI, USA), 10% v/v Hyclone FBS (Cytiva, Marlborough, MA, USA) and 1% v/v antibiotic-antimycotic (Thermo Fisher Scientific) at 37°C/5% CO_2_.

Working Cell Bank (MSC-WCB, Passage: 2) and Cell Therapy Product (MSC-CTP, Passage: 3) stocks of MSCs were used for the entirety of this study. Once thawed, cells were seeded at a density 3,000-3,500 cells/cm^2^ and weaned to xeno-free conditions using RoosterNourish-MSC (Rooster Bio, Fredrick, MD, USA). After 4 days, MSCs were dissociated from flasks using TrypLE Express Enzyme (Thermo Fisher Scientific) and counted using an NC-202 automated cell counter (ChemoMetec, Allerod, Denmark). Cell suspensions were centrifuged at 1100 rpm for 5 min and resuspended for encapsulation.

### Fabrication of microcapsules

Prior to encapsulation, cell suspensions were mixed with VitroGel-MSC (TheWell Biosciences, North Brunswick, NJ, USA) to a total volume of 6 mL in a 1:2 v/v ratio according to the manufacturer’s recommendations. This hydrogel precursor solution was loaded into a 10 mL syringe and mounted vertically onto a syringe pump (Harvard Apparatus, Halliston, MA, USA). Microcapsule generation was performed using a VARV1 Encapsulation Unit (Nisco Engineering AG, Zurich, Switzerland) at a voltage supply of 4.55 kV and 20 mL/h syringe pump flow rate. For all encapsulations a 28G nozzle supplied by Nisco Engineering AG was used and placed at a constant height of 3.2 cm from the collection basin. Electrosprayed microcapsules were allowed to crosslink for 4 hours in the collection basin containing 80 mL of RoosterNourish-MSC (Rooster Bio). After 4 hours, microcapsules were transferred to a PBS0.1 vertical-wheel bioreactor (PBS Biotech, Camarillo, CA, USA) and increased to a final volume of 90 mL. Bioreactors were maintained at an agitation rate of 25 rpm and 37°C. On day 3, a xeno-free RoosterReplenish-MSC-XF (Rooster Bio) was added at 2% v/v and the agitation rate was increased to 30 rpm.

### Characterization of microcapsules

Microcapsule density (capsules/mL) was acquired by electrospraying different volumes (1, 5, and 10 mL) of VitroGel-MSC at a consistent seeding density of 1.6×10^6^ cells/mL. The cell suspension to Vitrogel-MSC ratio was maintained at 1:2 v/v. After encapsulation, microcapsules were transferred to a PBS0.1 vertical-wheel bioreactor after 4 hours. The reactors were maintained at an agitation rate of 25 rpm and 37°C. On day 3, a xeno-free RoosterReplinish-MSC-XF (Rooster Bio) was added at 2% v/v and the agitation rate was increased to 30 rpm. On day 6, 1 mL samples were aliquoted to quantify the number of microcapsules using a Celigo Image Cytometer (Nexcelom Biosciences, Lawrence, MA, USA). Microcapsule size distribution was analyzed using ImageJ software.

### Taylor cone evaluation

Taylor cone formation was verified with a camera (Point Grey Research / FLIR, Grasshopper GS3-U3-41C6NIR-C (2048×2048 pixels, 5.5 μm pixel size) and four lenses (L1: Thorlabs AC-508-100-A-ML, L2: Thorlabs AC-508-100-A-ML, L3: Olympus Plan N, 4x / 0.1, L4: Thorlabs AC-254-150-A) to achieve appropriate magnification. Hydrogel precursor solution was loaded in a 10 mL syringe and the encapsulation unit was switched on to image the transition of droplets into a Taylor cone at an optimized voltage.

### Bioreactor sampling and metabolite analysis

Bioreactor sampling was performed to monitor cell growth kinetics, metabolite consumption, and waste accumulation throughout the culture time course. Microcapsules in suspension were sampled from bioreactors at 20 rpm agitation. A 3 mL sample was collected on day 1, while 1 mL samples were collected for remaining time points. Samples were incubated with CellTiter-Blue (Promega, Madison, WI, USA) at 20% sample volume according to manufacturer’s instructions for 4 hours. Data acquisition was followed as fluorescence readouts, made by the Varioskan LUX (Thermo Fisher Scientific) multimode reader. A multipoint reduction step was added in the SkanIt Software protocol session, and the average of the multipoint fluorescence signal for every well was calculated. After this step, a blank subtraction was carried out to account for any background reduction of resazurin occurring in control wells. Results were analyzed using a standard curve with linear regression analysis. Microcapsule free supernatants were analyzed using a Cedex Bioanalyzer (Roche Diagnostics, Indianapolis, IN, USA) for concentrations of glucose (mmol/L), lactate (mg/L), total protein (g/L), ammonia (mmol/L), lactate dehydrogenase (U/L), and glutamine (mmol/L). Microcapsules were stained with 2 μM Calcein AM (Thermo Fisher Scientific), and 4 μM Ethidium Homodimer-1 (Thermo Fisher Scientific) and imaged using a Zeiss Axio Observer. The following parameters were obtained from data acquisition:

Specific growth rate

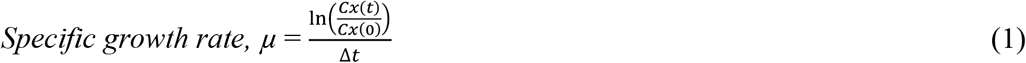

where µ is the specific growth rate (day^-1^), *C*_x_(t) and *C*_x_(0) are the final and initial cell numbers after time, *t* (days).

Population doublings

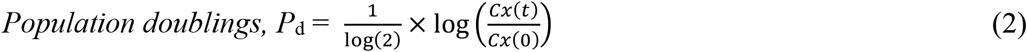

where *P*_d_ is the number of population doublings, and *C*_x_(t) and *C*_x_(0) are the final and initial cell numbers after time, *t* (days).

Specific metabolite consumption and waste production rate

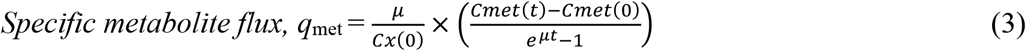

where *q*_met_ is the specific metabolite consumptions or waste production, µ is the specific growth rate (day^-1^), *C*_met_(t) and *C*_met_(0) are the final and initial metabolite concentrations, and C_x_(0) is the final cell number after time, *t* (days).

### Bioreactor harvest

Following a 6-day expansion in PBS0.1 vertical-wheel bioreactors, microcapsules were removed from the bioreactor and screened through a 40 μm nylon mesh cell strainer. Screened microcapsules were transferred back to bioreactors and resuspended in 60 mL dissociation solution that consisted of Cell Recovery Solution (TheWell Biosciences), 0.1% w/v L-Cysteine (Sigma-Aldrich), 0.1% v/v Phenol Red (Sigma-Aldrich), and NaOH to a pH of 7.0-7.5. 1 U/mL Papain (Sigma-Aldrich) was added to the bioreactor to initiate microcapsule degradation at 50 rpm for 30-45 mins. Cells suspensions were centrifuged at 1100 rpm for 5 min and counted using an NC-202 automated cell counter (ChemoMetec).

### Post bioreactor expansion

Cell health analysis of MSCs harvested from microcapsules was evaluated for their expansion capabilities. Cells were seeded at 200 cells/cm^2^ in Falcon T25 cm^2^ flasks (Corning Inc, Corning, NY, USA). Following a 7-day incubation, cells were dissociated from flasks using TrypLE Express Enzyme (Thermo Fisher Scientific) and counted using an NC-202 automated cell counter (ChemoMetec).

### Colony Forming Unit (CFU) assay

Hematopoietic stem cells (HSCs) were isolated from human bone marrow (Lonza) using the CD34 MicroBead Kit UltraPure (Miltenyi Biotech, Bergisch Gladbach, North Rhine-Westphalia, Germany), frozen, and stored at -180°C. 1×10^3^ HSCs and MSCs were resuspended in 0.1 mL Iscove’s Modified Dulbecco’s Medium, IMDM with 2% FBS (StemCell Technologies, Vancouver, BC, Canada). This HSC:MSC resuspension was added to 1 mL MethoCult H4034 Otimum (StemCell Technologies) and plated in a 6 well SmartDish (StemCell Technologies). CFU assays were quantified on day 14 using the STEMvision (StemCell Technologies) automated colony counter.

### Tri-lineage differentiation

Directed differentiation of MSCs into osteocytes, adipocytes, and chondrocytes was performed on microcapsules, and MSCs harvested from bioreactors on day 6. For osteogenic differentiation, MSCs were cultured in Mesenchymal Stem Cell Osteogenic Differentiation Medium (Sigma-Aldrich, Saint Louis, MO, USA) for 14 days, then fixed and stained with 2% Alizarin Red Stain Solution (Lifeline Cell Technology, San Diego, CA, USA). For adipogenic differentiation, MSCs were cultured in MesenCult Adipogenic Differentiation Kit (StemCell Technologies) for 14 days, then fixed and stained with Oil Red-O Solution (Sigma-Aldrich). For chondrogenic differentiation, MSCs were cultured in MesenCult-ACF Chondrogenic Differentiation Kit (StemCell Technologies) for 21 days, then fixed and stained with Alcian-Blue (Sigma-Aldrich). Phase contrast images were captured of stained differentiated and undifferentiated controls using a EVOS VL Core (Thermo Fisher Scientific).

### Lentiviral production

Lentiviral particles were produced using triple-transfection methods in adherent human embryonic kidney (HEK) 293T cells. Briefly, HEK293T cells were expanded in DMEM/F-12 media (Thermo Fisher Scientific) supplemented with 10% v/v FBS and 1% v/v antibiotic-antimycotic solution (Thermo Fisher Scientific). Cells were seeded at 40% confluency the day before transfection. HEK293T cells were co-transfected with pLV-EF1a-RFP (Vector Builder Inc, Chicago, IL, USA) and two packaging plasmids, psPAX2, plasmid #12260 (Addgene, Watertown, MA, USA) and pMD2.G, plasmid #12259 (Addgene), at a molar ratio of 3:2:1; and the transfection reagent polyethylenimine (Polyplus, New York, NY, USA) in OptiMEM (Thermo Fisher Scientific) medium for 15 min. DNA-PEI complex was then added dropwise to the cell culture. Transfection culture was carried out for 72 h until supernatant was collected, centrifuged, filtered through a 0.45 mm PES membrane filter and stored at -80°C. Vector titer was determined by qPCR using a Lentiviral titration kit (Applied Biologic Materials, Richmond, BC, Canada) on Quant Studio 3 (Thermo Fisher Scientific).

### Lentiviral transduction

Lentiviral particles were either added to a 2D MSC monolayer or to hydrogel precursor solution (3D model) at a multiplicity of infection (MOI) 50, in RoosterNourish-MSC medium (Rooster Bio). Selected groups received ViralEntry Transduction Enhancer reagent (Applied BiologicMaterials) at a ratio of 1:100 v/v. In 2D groups, media was replaced after 24 hours. In 3D groups, microcapsules were generated and harvested as previously described. Transduction efficiency was determined 10 days post-transduction via flow cytometry.

### Flow cytometry

Single cell suspensions of MSCs were stained for the following antibodies: CD34(MOPC-173) (BioLegend, San Diego, CA, USA), CD146(P1H12) (BioLegend), CD73(AD2) (Thermo Fisher Scientific), CD90(5E10) (Thermo Fisher Scientific) and CD105(SN6) (Thermo Fisher Scientific). Following surface marker staining, cells were fixed for 10 min in 2% paraformaldehyde and were analyzed using a FACS CANTO II (BD Biosciences, Franklin Lakes, NJ, USA). For lentiviral transduction efficiency assessment, cells were not stained, but directly fixed and analyzed using the same equipment. Data was analyzed using FlowJo software (BD Biosciences).

### Confocal microscopy

Z-stack images of microcapsules were stained with 5 μg/mL Hoechst 33342 (Thermo Fisher Scientific), 0.25 μM Calcein AM (Thermo Fisher Scientific), and 8 μM Ethidium Homodimer-1 (Thermo Fisher Scientific). Microcapsules were incubated at 37°C/5% CO_2_ for 30 minutes and imaged using a Zeiss 780 Confocal microscope.

## Results

### Evaluation of MSC growth in VitroGel-MSC and optimal seeding density in static culture

Initial studies determined the growth kinetics and expansion capability of MSCs cultured in static 2D Monolayer and 3D conditions using VitroGel-MSC at seeding densities ranging between 0.03125×10^6^ cells/mL and 0.50×10^6^ cells/mL (Fig. 1A). Cell density readouts were evaluated using CellTiter-Blue, which has been reported as a nontoxic, nondestructive metabolic approach to quantify cell dose in porous scaffolds [32]. Using optimized incubation conditions (Fig. S1), we determined that CellTiter-Blue is sensitive and reliable to quantify encapsuled MSCs cell densities without effecting hydrogel scaffold integrity. Figure 1A depicts the relationship between seeding density and growth kinetics of MSCs cultured in 2D Monolayer and 3D VitroGel-MSC conditions. This relationship demonstrates that MSCs cultured in 3D conditions have similar growth kinetics to their 2D counterparts and is further validated based upon MSCs grown in VitroGel-MSC achieving near equivalent or greater fold expansion compared to cells grown in 2D Monolayer (Fig. 1B). Notably, at densities of 0.25×10^6^ cells/mL and 0.50×10^6^ cells/mL, MSCs grown in 3D conditions had a significantly higher fold expansion, achieving a 2.0 and 1.625 greater expansion compared to their respective monolayer controls. In addition, there is an observed inverse correlation between growth rate and seeding density, with lower seeding densities yielding higher growth kinetics. VitroGel-MSC scaffold conditions were observed to significantly promote MSC expansion at all seeding densities over a six-day culture period albeit for 0.125×10^6^ cells/mL (Fig. 1C). Taken together, although a greater on average fold expansion was observed for seeding densities of 0.03125×10^6^ cells/mL and 0.0625×10^6^ cells/mL, these densities did not have statistical significance and were neglected from future studies.

**Fig. 1.**
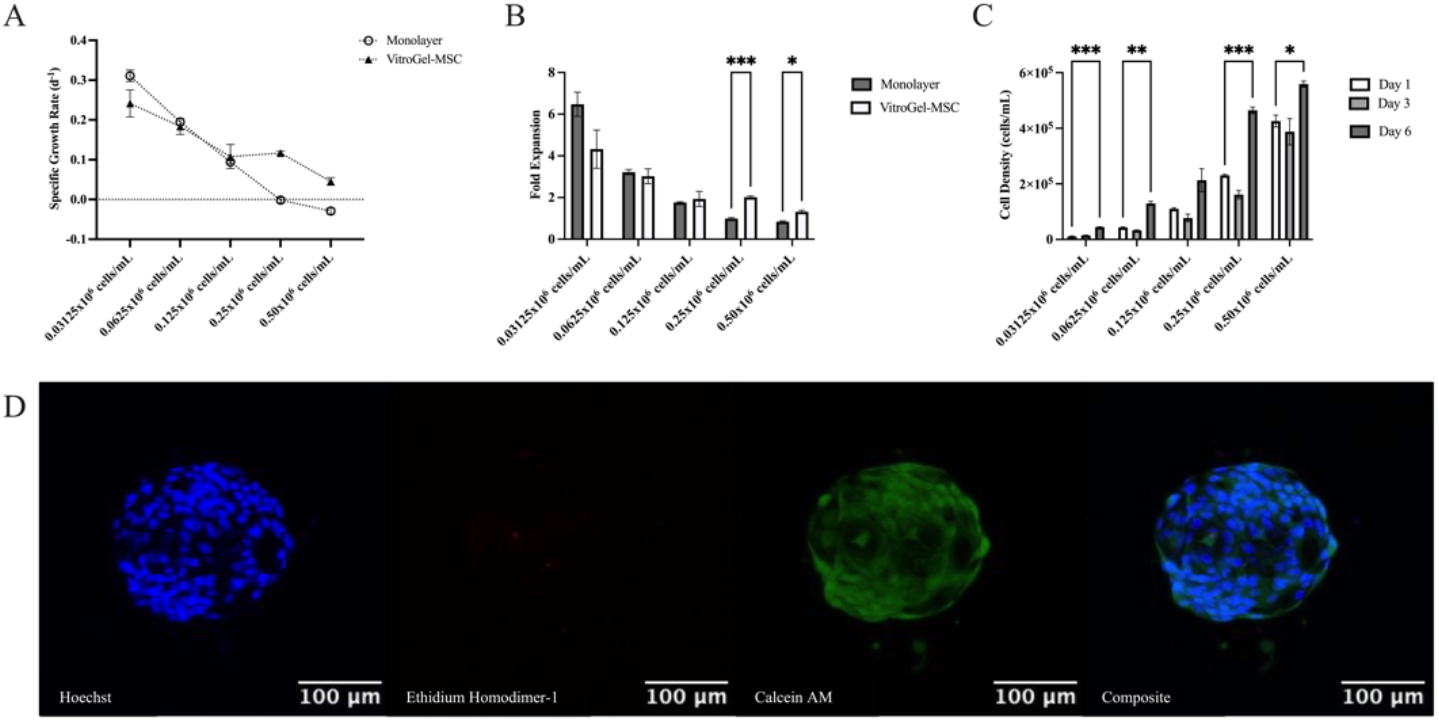
Evaluation of growth kinetics (A) and fold expansion (B) of MSCs grown in 2D Monolayer format vs. 3D VitroGel-MSC hydrogel system (n=3). (C) Time course expansion of MSCs cultured in VitroGel-MSC at varying seeding densities in static culture. Serial two-fold dilution of MSCs prepared in a 96-well plate (n=3). (D) Composite and single channel z-stack images of VitroGel-MSC capsules. Prior to imaging VitroGel-MSC capsules were stained with 0.5µg/mL Hoechst (Blue/Nuclei),0.25µM Calcein AM (Green/Live), and 8µM Ethidium Homodimer-1 (Red/Dead).

To visualize the 3D growth of MSCs in VitroGel-MSC, we seeded MSCs within the hydrogel at a density of 0.25×10^6^ cells/mL and created cell laden droplets in a standard well plate. After a six-day culture, we observed well defined cell laden capsules (Fig. 1D). Z-stack confocal imaging indicated the presence of viable MSC nuclei with minimal cell death, and encapsulated cells assuming natural spindle morphology. These results suggest that VitroGel-MSC can provide encapsulated cells with a tailored 3D microenvironment conducive for MSC proliferation and it is feasible to create cell laden capsules to be translated into a vertical-wheel bioreactor system.

### Electrospraying and characterization of MSC microcapsules

A small-scale electrospraying system was designed to automate and increase production of MSC microcapsules. Figure 2A depicts a cell-polymer solution loaded into a syringe that is extruded under an electric field to form cell microcapsules that polymerize once exposed to a bath of cell culture medium. Liquid atomization, a phenomenon where the electric field overcomes the surface tension of the liquid, is a decisive factor in achieving a narrow microcapsule size distribution. To determine the critical voltage of the liquid, wherein the droplets (Fig. 2B) transition into a steady stream of uniform droplets; a high precision imaging technique was used. Taylor cone (Fig. 2C) was observed at a voltage of 4.55kV (Vid. S1). This voltage was used for the remainder of studies.

**Fig. 2.**
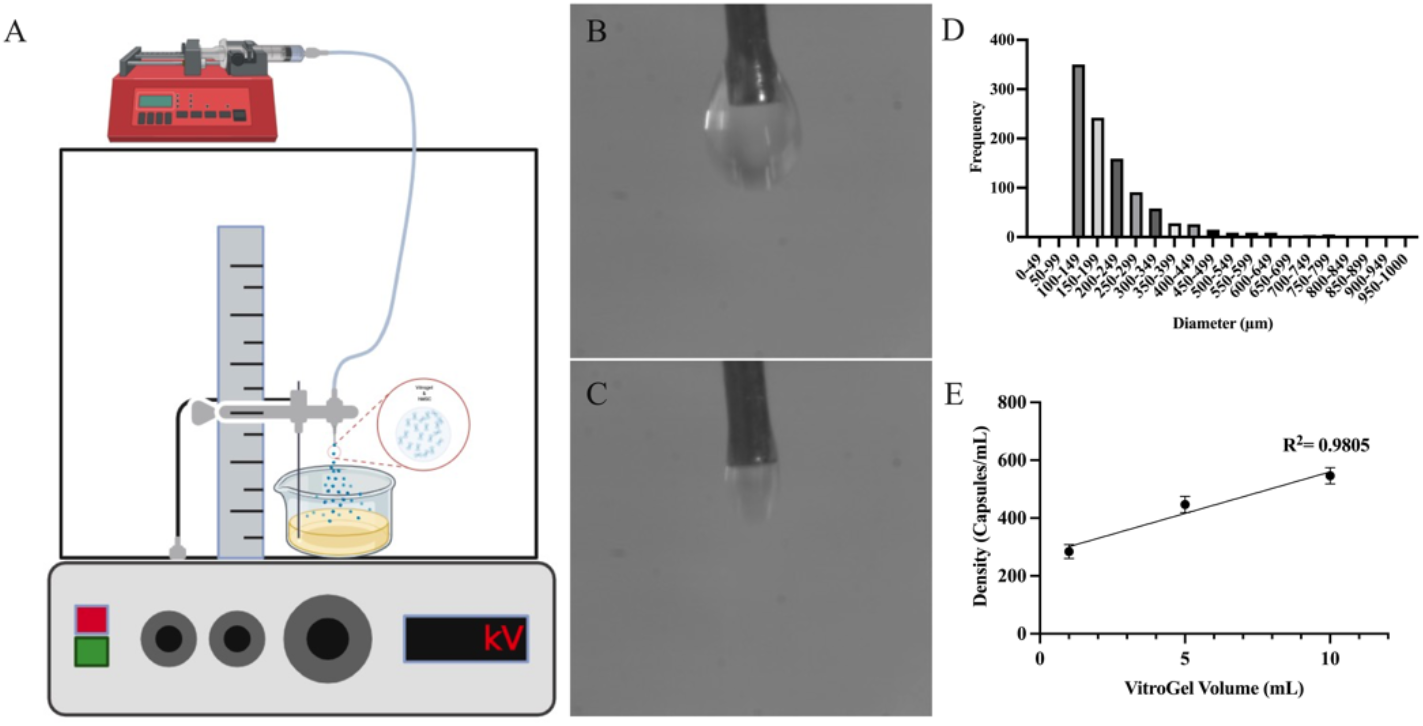
(A) Schematic of VitroGel-MSC microcapsule formation using the Nisco VAR-VI encapsulator. (B-C) Representative images capturing a transition from individual to steady stream droplets, indicated by Taylor cone formation. (D) Size distribution and frequency of VitroGel-MSC microcapsules formed (n>1000). (E) Linear correlation between VitroGel-MSC volume and microcapsule output.

Microcapsule size manufactured per batch is designated as a validation test for the repeatability of this encapsulation platform. An ideal biomanufacturing technique would produce microcapsules of consistent size irrespective of the VitroGel-MSC volume used, provided that cell seeding density is fixed. An analysis of size distribution (Fig. 2D) confirmed that manufactured microcapsules (n>1000) exhibit a uniform size with a higher frequency of microcapsules ranging between 100-149 µm (34.7%) followed by 150-200 µm (24%), and few microcapsules ranging between 400-500 µm (4.06%)

The volume of VitroGel-MSC used in the biomanufacturing of MSCs is a key input parameter of the electrospraying process with a resultant output of number of cell-laden microcapsules. To establish the relationship of hydrogel volume and microcapsule output, MSC encapsulation experiments (n=3) with different VitroGel-MSC volumes were conducted at a consistent seeding density of 1.6×10^6^ cells/mL. The study found that the number of microcapsules produced per encapsulation were correlated to the volume of electrosprayed VitroGel-MSC. For this specific small-scale batch size, 10 mL of Vitrogel-MSC produced 546 ± 28 microcapsules, 5 mL of Vitrogel-MSC produced 447 ± 28 microcapsules, and 1 mL of Vitrogel-MSC produced 228 ± 24 microcapsules. The plotted graph (Fig. 2E) shows a linear correlation with a fit of R^2^ =0.98.

### MSC expansion in vertical-wheel bioreactors utilizing a fed-batch process

Using a fixed and optimized set of input parameters, we developed a biomanufacturing workflow (Fig. 3A) to evaluate expansion of MSCs in a fed-batch process at seeding densities of 0.125×10^6^ cells/mL, 0.25×10^6^ cells/mL, and 0.50×10^6^ cells/mL. Encapsulation efficiency was evaluated as the viable cell number twenty-four hours post encapsulation relative to the total number of cells encapsulated on day zero (Fig. 3B). VitroGel-MSC microcapsules seeded at a density of 0.125×10^6^ cells/mL had the highest encapsulation efficiency of 127% ± 11%, compared to encapsulation efficiencies of 72% ± 6%, and 83% ± 5% for densities of 0.25×10^6^ cells/mL and 0.50×10^6^ cells/mL respectively.

**Fig. 3.**
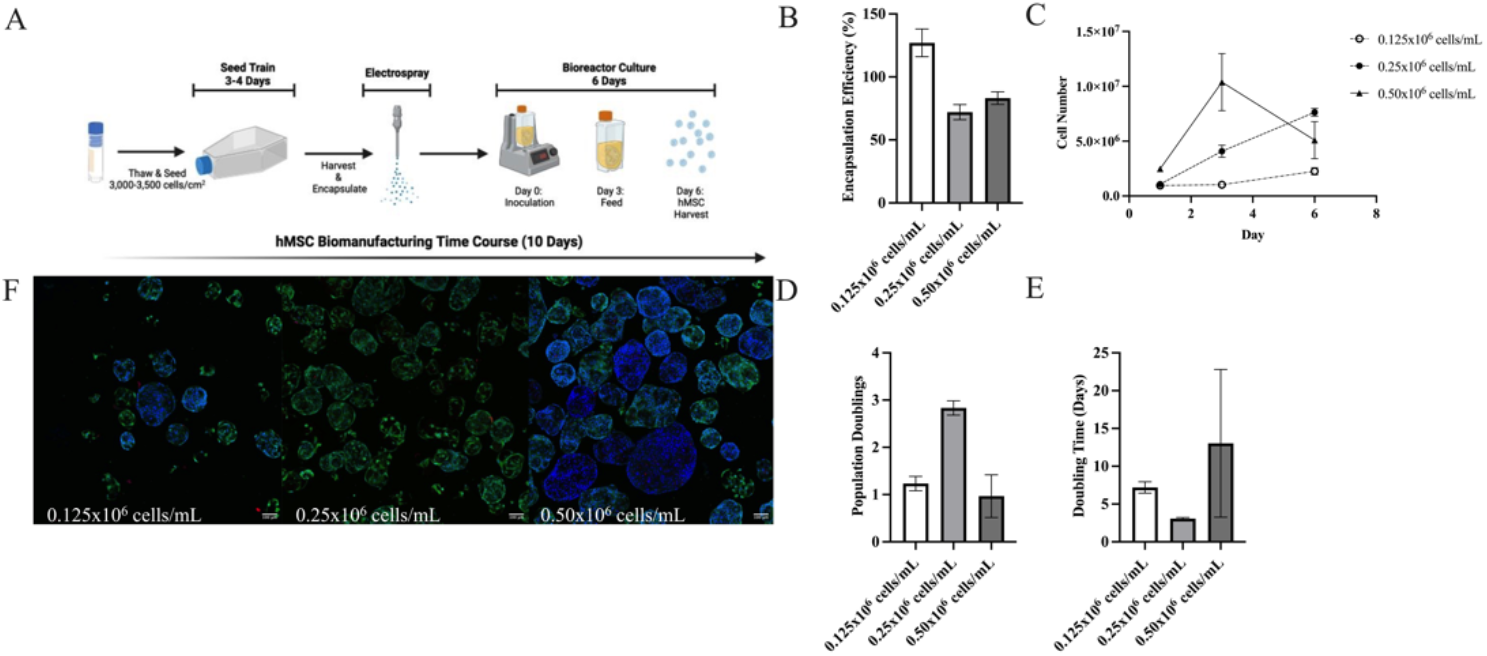
(A) Schematic of MSC biomanufacturing time course. (B) Encapsulation efficiency (C) growth profile (D) population doublings and (E) doubling time of VitroGel-MSC microcapsules at varying encapsulation densities (n=3). (F) Composite Z-stack images ofVitroGel-MSC microcapsules at varying encapsulation densities. Prior to imaging VitroGel-MSC microcapsules were stained with 0.5µg/mL Hoechst (Blue/Nuclei), 0.25µM Calcein AM (Green/Live). and 8µM Ethidium Homodimer-1 (Red/Dead).

The growth profile of cell-laden VitroGel-MSC microcapsules maintained in suspension culture were evaluated over a six-day time course with a 2% v/v feed on day 3. Incorporating a day 3 feed into the biomanufacturing time course eliminates the need for media exchanges that otherwise would be costly in larger bioreactor systems. Day 6 cell counts showed yields of 2.25×10^6^ ± 3.21×10^5^ cells (∼ 2.36-fold expansion), a yield of 7.63×10^6^ ± 3.46×10^5^ cells (∼ 7.04-fold expansion), and a yield of 5.10×10^6^ ± 1.69×10^6^ cells (∼ 2.56-fold expansion) for microcapsules electrosprayed at encapsulation densities of 0.125×10^6^ cells/mL, 0.25×10^6^ cells/mL, and 0.50×10^6^ cells/mL respectively (Fig. 3C). Among the three encapsulation densities, microcapsules electrosprayed at a density of 0.50×10^6^ cells/mL achieved the highest cell yield of 10.38×10^6^ ± 2.60×10^6^ cells (∼ 4.20-fold expansion), however plateaued beyond day 3. Similar proliferation trends were observed in each encapsulation density’s population doubling (Fig. 3D) and MSC doubling time (Fig. 3E). MSCs encapsulated at a density of 0.25×10^6^ cells/mL achieved the highest population doubling (2.82 ± 0.15), with the lowest doubling time of 3.1 ± 0.16 days. Notably, after six days of expansion, there is an absence of microcapsule aggregation amongst all densities (Fig. 3F & Fig. S2 respectively).

Metabolite consumption, waste production, and the net metabolite flux were monitored to better understand potential factors affecting the expansion process (Fig. 4). Medium analysis indicated a consistent metabolic consumption of glucose and glutamine amongst all encapsulation densities. Interestingly, when evaluating the net metabolite flux per encapsulation density over the expansion time course, encapsulations at 0.125×10^6^ cells/mL demonstrated a preference for glutamine consumption. Between days 0-1, the rate of glutamine consumption was 0.24 ± 0.11 pmolcell^-1^d^-1^ and sharply increased between days 1-3 and 3-6 to rates of 1.67 ± 0.09 pmolcell^-1^d^-1^ and 2.06 ± 0.00 pmolcell^-1^d^-1^ respectively. This increased rate of glutamine consumption was observed to coincide with higher rates of ammonia (NH3), LDH, and Total Protein production during the same time phase. 0.25×10^6^ cells/mL and 0.50×10^6^ cells/mL encapsulation densities demonstrated a preference for glucose consumption, which coincided with rates of lactate waste production. Between days 1-3 and 3-6, the glucose consumption rate for 0.50×10^6^ cells/mL encapsulations sharply increased from 2.04 ± 0.03 pmolcell^-1^d^-1^ to 7.00 ± 1.35 pmolcell^-1^d^-1^, and the lactate production rate increased from 387 ± 43.7 pgcell^-1^d^-1^ to 1090 ± 113.8 pgcell^-1^d^-1^. Increased ammonia (NH3), LDH, and Total Protein production rates were observed, however, were not as high as 0.125×10^6^ cells/mL metabolite rates.

**Fig. 4.**
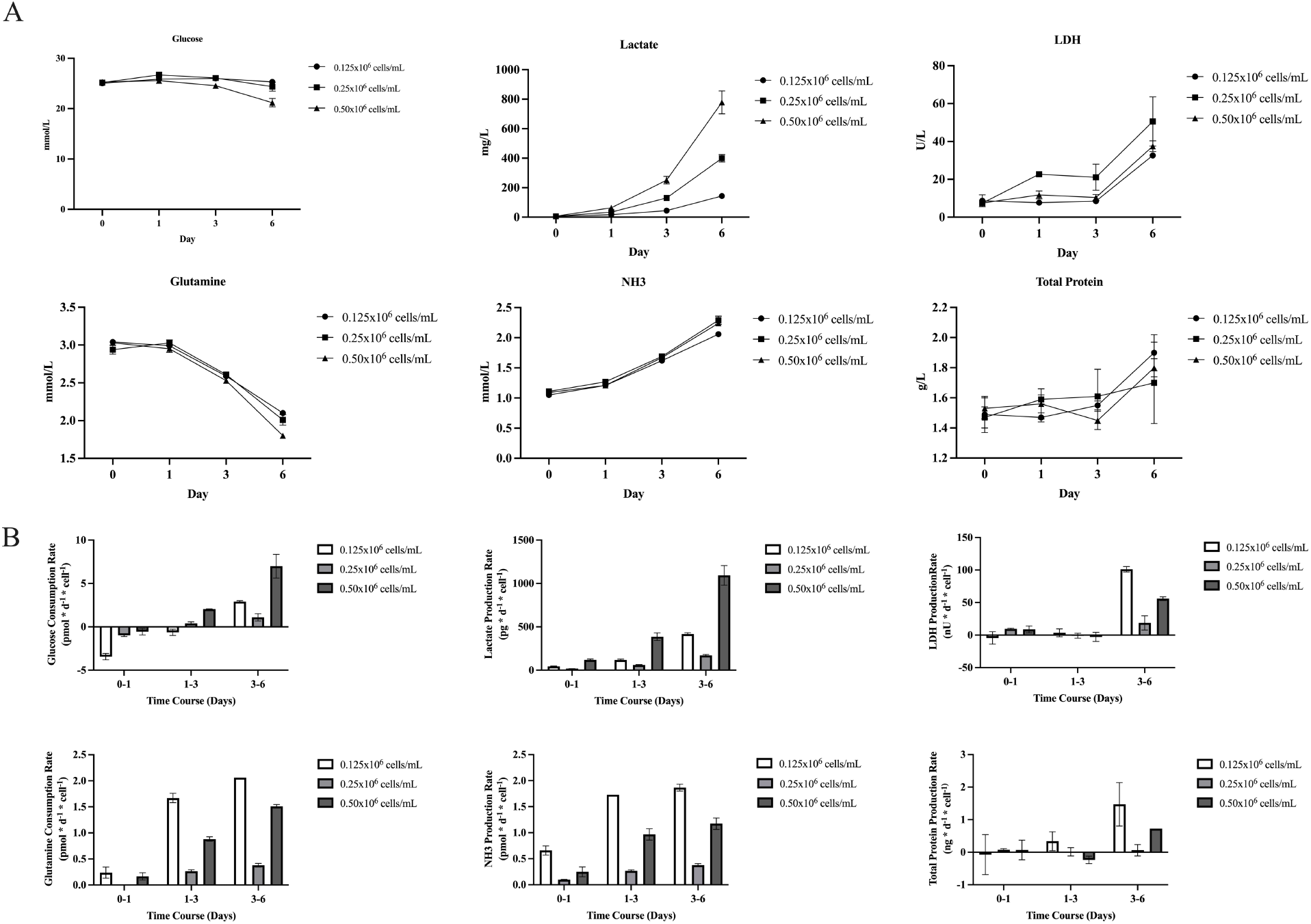
(A) Metabolite consumption and waste production. (B) Net metabolite flux per cell for MSCs expanded in VitroGel-MSC microcapsules over a 6-day time course. Microcapsules were maintained in suspension culture in a vertical-wheel bioreactor at an agitation rate of 25 rpm. On day 3, the agitation rate was increased to 30 rpm.

### Microcapsule digestion and evaluation of MSC Critical Quality Attributes (CQAs)

Following expansion in vertical-wheel bioreactors, MSCs encapsulated at a density of 0.25×10^6^ cells/mL were harvested at their peak cell yield on day 6. The choice of dissociation solution has implications that can affect harvest yield, cell viability, and MSC critical quality attributes which define the formulated and filled product from a manufacturing perspective. Here, we evaluated the choice of dissociation solution to degrade VitroGel-MSC microcapsules and reconstitute MSCs into single-cell suspension. We screened dissociative solutions that have been previously reported with MSCs and VitroGel platforms, and include Accutase, Cell Recovery Solution, TrypLE Express, and Papain (Fig. 5A). Despite being a non-enzymatic cell harvesting solution frequented with VitroGel platforms [33, 34] Cell Recovery Solution alone was unable to reconstitute MSCs into single-cell suspension potentially due to encapsulated MSCs achieving a high density and extracellular network within VitroGel-MSC microcapsules. After 30 min of digestion, 83% ± 7% of VitroGel-MSC microcapsules remained non-degraded (Fig. 5B). Similarly, after 30 min of digestion, 86% ± 5% and 100% ± 0% of VitroGel-MSC microcapsules remained non-degraded for Accutase and TrypLE Express dissociative solutions. Papain (1 U/mL) was observed to significantly digest VitroGel-MSC microcapsules, with 15% ± 11% remaining non-degraded and harvested MSCs maintaining 94.3% ± 0.82% viability. With MSCs isolated from sampled microcapsules on day 6, we performed a general panel of critical quality attributes (CQAs) to determine if MSCs maintained their attributes and functionality following suspension culture in a vertical-wheel bioreactor. Cells harvested from vertical-wheel bioreactors (MSC-BIO) were evaluated for cell health, functionality, surface marker expression, and tri-lineage differentiation potential in comparison to cell bank stocks (MSC-WCB and MSC-CTP).

**Fig. 5.**
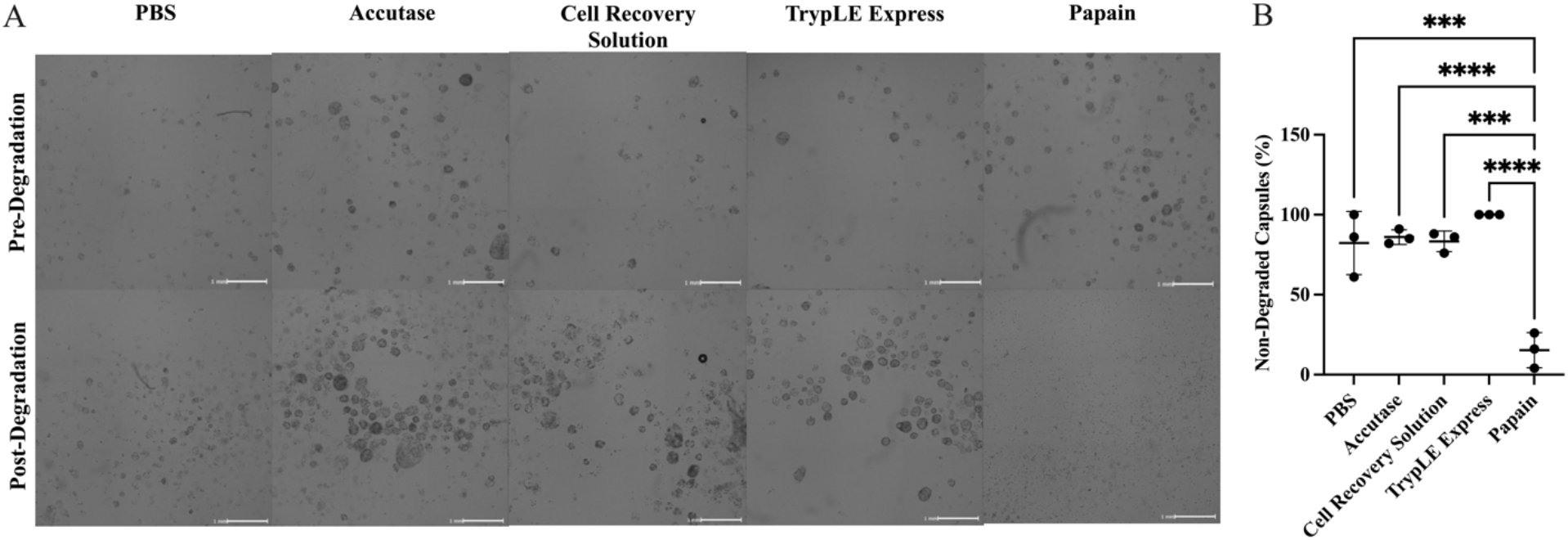
(A) Well images of VitroGel-MSC microcapsules pre- and post-enzymatic degradation (n=3). Microcapsules were allowed to digest for 30 min at 37°C. Data analysis and images were analyzed using Celigo Image Cytometer. (B) Percent of capsules remaining after 30 min of enzymatic degradation(n=3).

As an indication of cell health, we compared the proliferative capabilities of MSC-WCB (Passage 2), MSC-CTP (Passage 3), and MSC-BIO in 2D monolayer conditions over a 14-day culture. All MSC passages maintained their characteristic spindle morphology, however, MSC-WCB stocks achieved a significantly greater on average cell density of 4.74×10^5^ ± 5.63×10^4^ cells/mL compared to average cell densities of 2.90×10^5^ ± 3.89×10^4^ cells/mL, and 1.73×10^5^ ± 3.82×10^4^ cells/mL for MSC-CTP and MSC-BIO respectively (Fig. 6A). Metabolic consumption and waste production also exhibited similar trends with MSC-WCB stocks consuming and producing more glucose and lactate than either MSC-CTP or MSC-BIO.

**Fig. 6.**
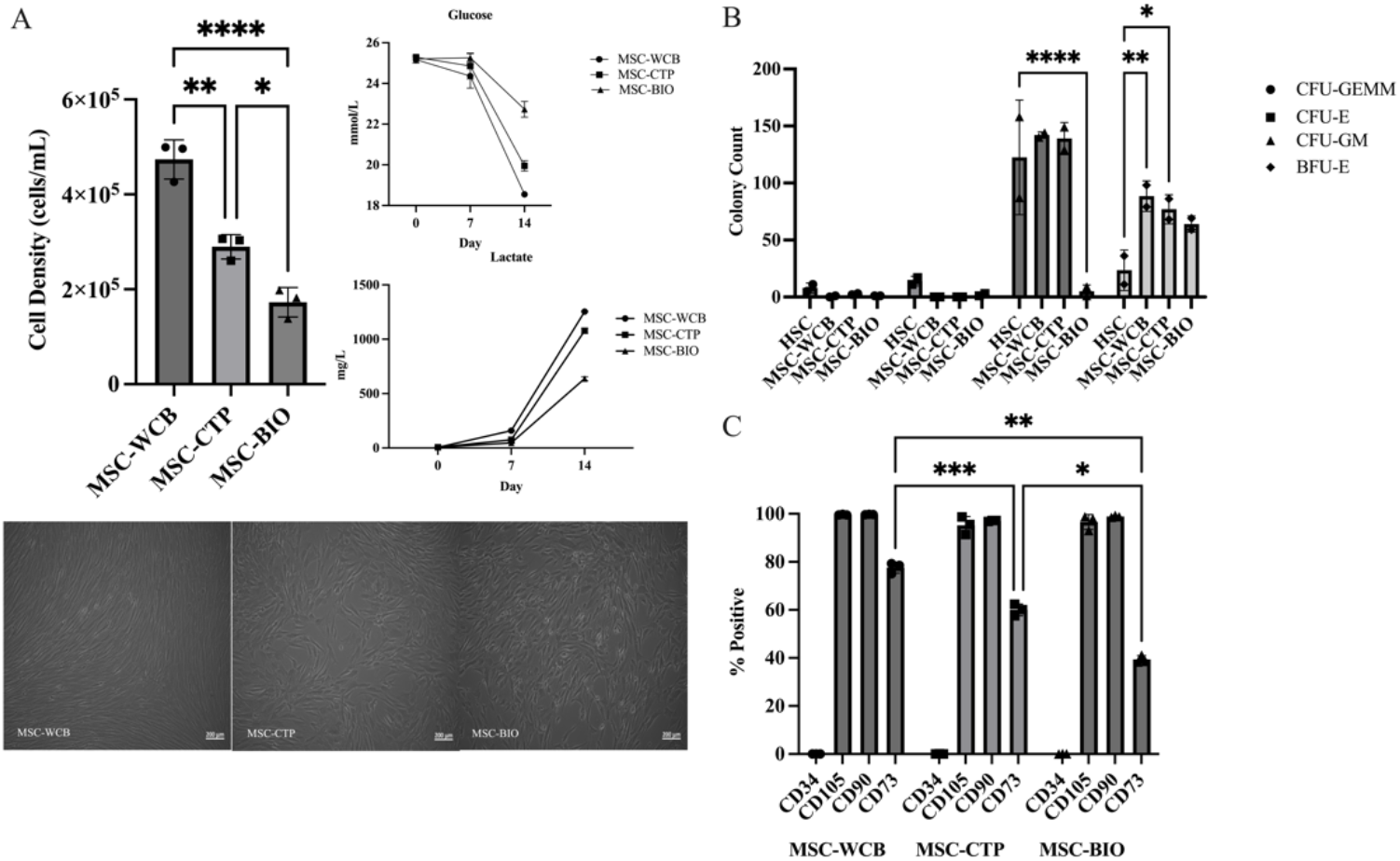
MSC characterization following suspension culture in a vertical-wheel bioreactor. (A) Post bioreactor expansion, metabolite consumption, waste production, and cell morphology evaluation as a validation of cell health. (B) Day 14 Colony Forming Unit (CFU) assay evaluating MSC signaling functionality (C) Evaluation of MSC cell surface marker expression.

Cell functionality was evaluated based upon how MSCs influenced the differentiation of human hematopoietic stem cells (HSCs) into progenitor cell populations (Fig. 6B). HSCs co-cultured with MSC-BIO maintained similar commitments toward colony forming units (CFU) of granulocyte, erythrocyte, macrophage, megakaryocyte (CFU-GEMM) and erythroid (CFU-E) progenitors. Notably, HSCs co-cultured with MSC-BIO resulted in significantly lower granulocyte, macrophage (CFU-GM) progenitors. All MSC passages influenced a greater HSC commitment toward burst forming unit erythroid (BFU-E) progenitor colonies, however only HSCs co-cultured with MSC-WCB and MSC-CTP were statistically significant. We also observed MSC-BIO maintained CQAs of surface marker expression for CD34, CD105, and CD90. Interestingly, CD73 expression significantly decreased (∼20%) with each successive passage (Fig. 6C). Based upon a significant difference in CD73 expression within cell stocks MSC-WCB (77.6%) and MSC-CTP (60.0%), representing increasing accrual of population doublings it suggests that CD73 may be a useful marker of cell age. In addition, MSC-BIO maintained osteogenic, adipogenic, and chondrogenic differentiation potential from harvested cell populations and even *in situ* if differentiation conditions were applied to cellular microcapsules (Fig. 7 & S4, respectively).

**Fig. 7.**
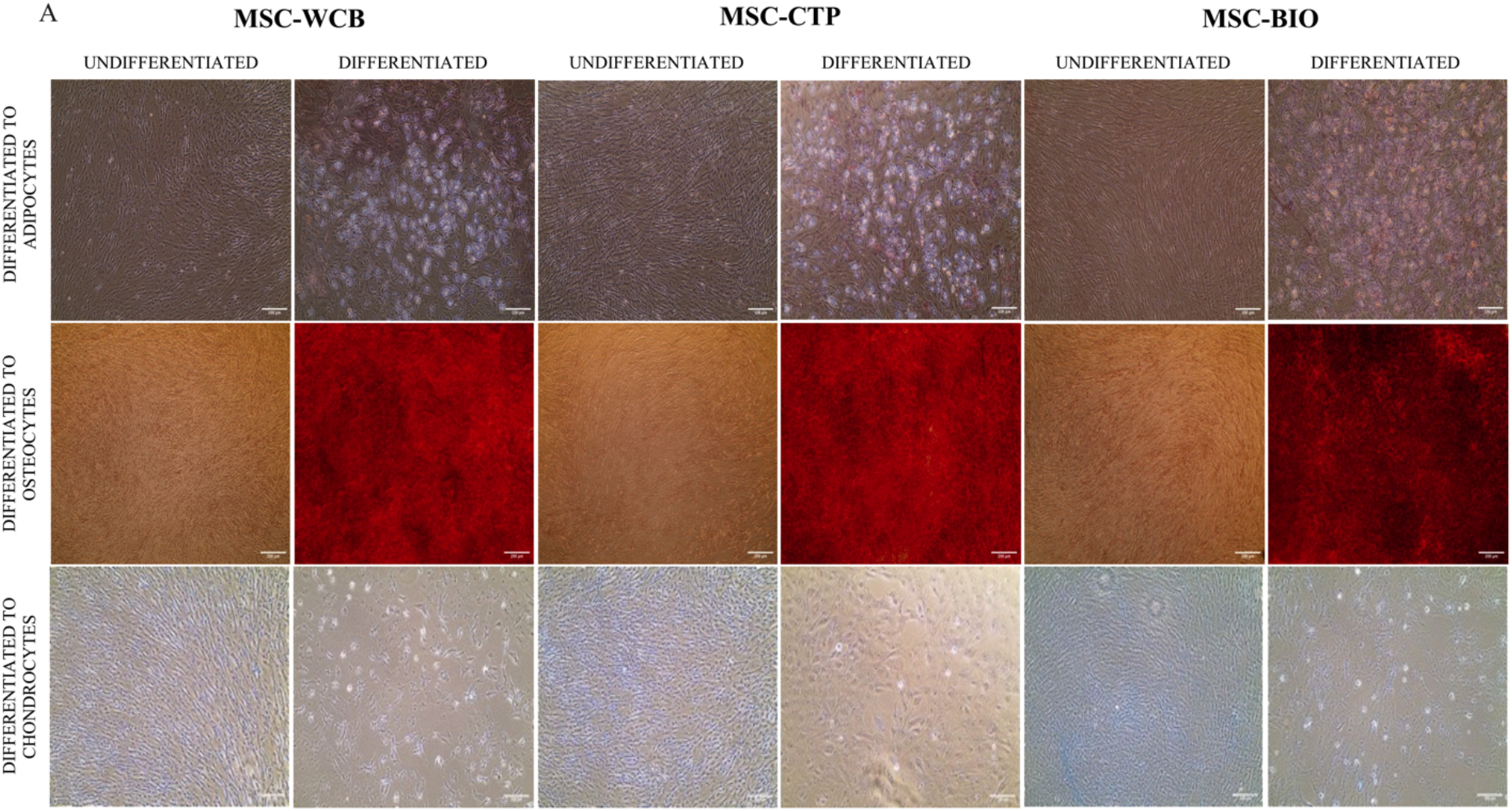
(A) Tri-lineage differentiation potential of cell bank stocks (MSC-WCB & MSC-CTP), and cells harvested from VitroGel-MSC microcapsules into adipocytes, osteocytes, and chondroycytes.

### Genetic manipulation of MSCs within VitroGel-MSC microcapsules

Viral vectors have been increasingly explored as gene delivery tools for induction of long-term transgene expression in MSCs, widening the potential of these cells as gene therapy agents [35]. Ranging from cardiac regeneration [36, 37] to targeted treatment of bone defects [38], autoimmune disorders [39] and cancer applications [40], genetic manipulation of MSCs has proven to be a promising tool in tissue engineering and regenerative medicine. Herein, we sought to demonstrate the functionality of encapsulated MSCs by employing our biomanufacturing system to promote lentiviral transduction. As a proof-of-concept, VitroGel-MSC microcapsules were co-electrosprayed with lentiviral vectors constitutively expressing a red fluorescent protein (RFP). RFP expression within VitroGel-MSC microcapsules was detected on day 6 (Fig. S5), however transduction efficiency was not assessed until day 10 and compared to a 2D monolayer transduction control. Remarkably, our 3D model was able to recapitulate about 60% of standard monolayer transduction efficiencies (Fig. 8A). Importantly, the use of a viral entry enhancer reagent was crucial to achieve such levels of transduction, as shown in Fig. 8B. Images of genetically modified VitroGel-MSC microcapsules at day 10 post lentiviral transduction confirm stable RFP expression (Fig. 8C), demonstrating successful genetic manipulation of MSCs under our 3D biomanufacturing conditions.

**Fig. 8.**
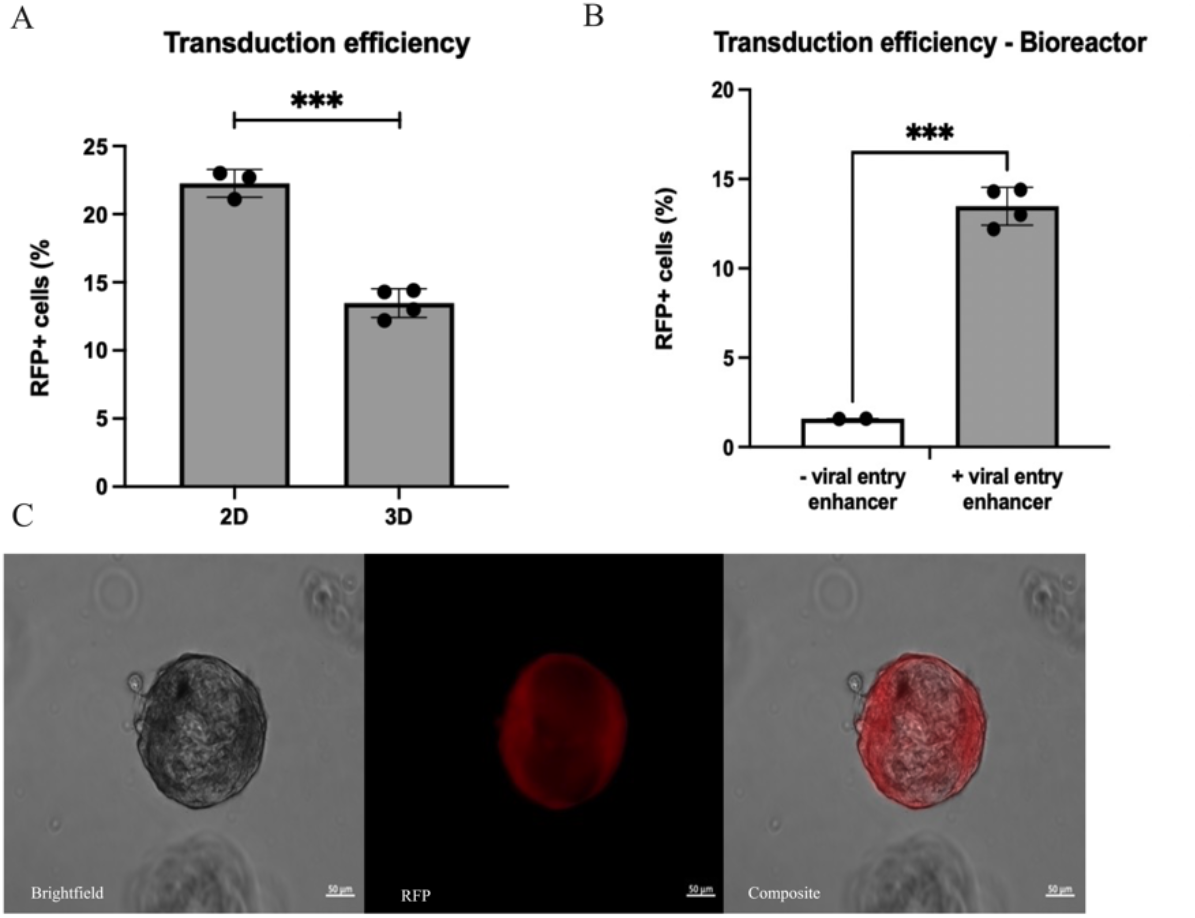
Lentiviral transduction efficiency of VitroGel-MSC microcapsules within vertical-wheel bioreactors (A) in comparison to 2D monolayer controls (n=2) (B) with (n=2) and without (n=l)a viral entry enhancer. (C) Day 10 single channel and composite images of VitroGel-MSC microcapsules co-electrosprayed with lentiviral vectors constitutively expressing a red fluorescent protein (RFP).

## Discussion

MSC based therapeutic for adults require consideration to be manufactured to meet clinical demand. Various approaches and process optimizations have been investigated to transition from adherent 2D to suspension-based 3D cultures for the commercial production of MSC therapeutics, with several research initiatives focusing primarily on microcarrier [41-43] or microencapsulation modes of expansion [28, 44, 45].

The results herein identify a proof-of-concept manufacturing platform that presents as a hybrid solution to expand MSCs and potentially other adherent cell types of interest to the biopharmaceutical sector. Prior MSC microencapsulation initiatives that utilize both synthetic and natural hydrogels have demonstrated limited expansion capability or utilize encapsulation protocols that are difficult to scale at commercial production. Kumar *et al*. [26] utilizes an electrospraying platform to encapsulate MSCs within alginate-based capsules. These capsules were cultured both *in vitro* and *ex vivo*, however over a seven-day time course cell viability was reduced by fifty percent. Perera *et al*. [46] photopolymerizes a vortex-induced emulsion of hydrogel precursor solution (<1 mL) to encapsulate MSCs within PEGDA microspheres, however this approach to microencapsulation is limited in scale-out, cost ineffective, and susceptible to inter-operator variability. We instead have opted to evaluate a xeno-free, polysaccharide hydrogel, VitroGel, that is commercially available for 3D expansion of MSCs to enable wide access to a quality-controlled material for community use. Within this study, we have studied several process parameters associated with electrosprayed based encapsulations of MSCs. To our knowledge, never-before has this hydrogel been subjected to electrospraying, to encapsulate MSCs within microcapsules for bioprocess engineering. Initial cell proliferation studies in static culture have shown that MSCs cultured in 3D conditions using VitroGel-MSC have similar growth kinetics to MSCs grown in 2D monolayer at varying densities. Aside from comparing growth kinetics we have optimized the encapsulation density to yield optimal MSC expansion potential. It is worth noting that the MSCs used in the entirety of this study were derived from a single donor. Several studies have suggested inter-donor variance can influence MSC expansion potential and functionality [47, 48]. From this perspective to avoid heterogeneous outcomes, future studies should evaluate MSC expansion potential across multiple donors prior to encapsulation. In addition, VitroGel matrices are also available in several variations, of which future work can screen for optimized MSC growth profiles and manipulate polymerization kinetics.

One process parameter to set when electrospraying is the applied voltage. The effects of applied voltage to produce a single cone jet mode characterized by Taylor cone formation during electrospraying remains contested. An applied voltage should produce a steady stream of microcapsules without affecting cell viability or inducing needle vibration. Gryshkov *et al*. [49] reports applied voltages (15-25 kV) do not hamper the viability of encapsulated MSCs within alginate beads. Whereas Qayyum *et al*. [50] in contrast has reported MSCs encapsulated in electrosprayed PEG microspheres have a significantly reduced viability (< 70%) at an applied voltage of 15 kV. Our results indicate a critical voltage of 4.55 kV was sufficient to produce a steady stream of VitroGel-MSC microcapsules, well below reported voltages that would impact cell viability. The encapsulation of cells within VitroGel-MSC also achieved high encapsulation efficiency, as scale-up with low efficiency can lead to an increase in the cost of manufacturing. Amongst all densities, the range of encapsulation efficiency quantified twenty-four hours after encapsulation was observed between 64% - 142% with an average of 94%, which exceeds microcarrier based cell attachment efficiencies [51] ranging between 42% - 142% with an average of 84%.

A major limitation of microcarrier based modes of expansion is aggregation leading to cell-detachment during the expansion process. Such limitations have required manufacturers to either supplement additional microcarriers into the bioreactor thereby increasing the available surface area, increasing the agitation rate, addition of detergents or combinations thereof; however, both risk impacting the MSC functionality and phenotype [23, 52]. While evaluating the effect of microcarrier aggregation on cell growth Lam *et al*. [53] concluded that microcarrier aggregates between 200-400 µm are conducive for the expansion of cells, however, higher agitation rates were required for cell detachment. Such high agitation rates during cell recovery can be detrimental for cell viability [54]. VitroGel-based encapsulation can help rectify manufacturing constraints imposed by microcarrier aggregation. More than fifty percent of our manufactured VitroGel-MSC microcapsules have ranged in size between 100-200 µm and had no observable aggregation after six days of expansion (Fig S2). Few public reports elucidate bead generators that can be scaled to commercial production. From the reports that are available, high-throughput production of cell-laden capsules has been achieved using a multi-nozzle extrusion head [55]. Notably, Swioklo *et al*. [56] utilizes an extrusion head containing nine nozzles to produce alginate beads in a drop-wise method at a rate of 3500 beads per minute. Adapting a similar approach to our platform whereby VitroGel-MSC microcapsules are produced at a higher rate in the presence of an electric field may pose a potential scale-up solution.

Variations in MSC growth within microcapsules depicted in Fig. 3 shows the capabilities and limitations of this system that should be considered to ensure optimal expansion potential in future scaled-up runs. Optimal yields at an encapsulation density of 0.25×10^6^ cells/mL achieved an ∼7x expansion of encapsulated MSCs within six days. Based upon specific metabolite rates that were monitored throughout the expansion time course, we believe the performance of 0.25×10^6^ cells/mL microcapsules is attributed to more permissible metabolite consumption and waste production levels.

Metabolite flux results for encapsulations at a density of 0.125×10^6^ cells/mL suggest waste accumulation affected expansion potential. Interestingly, trends in net metabolite flux for 0.125×10^6^ cells/mL microcapsules showed significant glutamine consumption, which coincided with sharply higher productions rates of ammonia. To monitor cell death, we monitored LDH during expansion as an indirect measure of lysed cells into culture supernatant. Despite having the lowest encapsulation density, we observed the highest rates of LDH. These results would suggest the expansion potential of 0.125×10^6^ cells/mL microcapsules was affected by cytotoxic levels of ammonia waste that resulted in cell lysis. Similar trends have been reported by Schop *et al*. [57] who observed ammonia and lactate accumulation inhibited cell growth once concentrations of 2.4 mM ammonia and 35.4 mM of lactate was achieved.

The expansion potential of 0.50×10^6^ cells/mL encapsulation instead suggested this system reaches a spatial capacity due to a limited availability of surface area within VitroGel-MSC microcapsules. Growth trends as reported in Fig. 3 indicate that MSCs within 0.50×10^6^ cells/mL microcapsules reach a peak yield on day 3 and plateau until final read outs on day 6. Since we used a fed-batch process that integrated a re-feed on day 3, it is unlikely that a deprivation of media nutrients would have contributed to this effect. This plateau in cell yield between days 3-6 may instead be attributed to increased rates of lactate production that inhibited cell growth. Although ammonia, LDH and Total Protein production rates increased, their rates of production were not observably high compared to 0.125×10^6^ cells/mL encapsulations. Notably, we can ascertain a linear relationship between microcapsule output and VitroGel volume, however, the effective surface area within each microcapsule remains ambiguous, of which maybe a crucial parameter for scaled-up commercial runs. Further investigation to quantify available surface area will not only provide insight into observed growth trends within microcapsules but will also provide a standardization to monolayer or microcarrier based cell expansion.

A successful biomanufacturing platform implies expansion of the target population without loss in functionality and immunophenotype of the cell. Notably, downstream processing of cell therapy products requires significant optimization as the process can affect cell viability and functionality [58]. Most downstream processing involved in biologics is designed to isolate byproducts of expansion such as proteins, antibodies, or cell secretions as exosomes without recovering cells as the desired product [59]. Therefore, it is necessary to develop a robust harvest protocol, which has minimal effect on the quality parameters. To ensure maximum cell recovery, we screened multiple disassociation agents and identified papain, as an enzymatic dissociative capable of reconstituting cells into single cell suspension. Not commonly used in traditional cell culture, papain has been used for the disassociation of human MSC aggregates [60] and used to digest CNS tumors into a single cell suspension [61]. It is also worth noting that encapsulations for 0.125×10^6^ cells/mL and 0.50×10^6^ cells/mL densities were conducted with MSC-CTP (Passage 3) cell bank stocks, whereas encapsulations for a density of 0.25×10^6^ cells/mL were conducted with MSC-WCB (Passage 2) cell bank stocks. MSC characterization results whereby the expansion potential of MSC-WCB and MSC-CTP cell bank stocks was compared to the expansion potential of MSCs harvested from vertical-wheel bioreactors, MSC-BIO (Passage 3), indicated a significant decrease correlated with cell passage. This significant difference presents itself as a limitation to this study as it could have affected the performance of 0.125×10^6^ cells/mL and 0.50×10^6^ cells/mL density encapsulations. Since MSC-CTP and MSC-BIO cells are of the same passage, results would also suggest a decrease in proliferation can be attributed either to conditions within vertical-wheel bioreactors or due to microcapsule processing with papain. Although one concentration of papain was used for the entirety of this study and was found to maintain >90% viability, additional investigation into various concentrations of papain and its effect on the maintenance of critical quality attributes should be evaluated.

Finally, we evaluated the use of our 3D bioreactor platform for concomitant viral transduction and expansion of MSCs. Genetically engineered MSCs have far-reaching potential as therapeutics, with a broad scope of applications. Both viral and non-viral gene delivery approaches have been extensively investigated in the fields of tissue engineering, regeneration and oncology using MSCs [62-64]. Mangi *et al*. overexpressed the prosurvival gene Akt1 in MSCs using lentiviral vectors, successfully repairing infarcted myocardia and restoring cardiac performance [37]. Zhu and group were able to suppress the growth of gastric cancer xenografts by treating mice with genetically engineered MSCs overexpressing NK4, an antagonist of hepatocyte growth factor receptors [40]. Andrews *et al*. genetically engineered MSCs with recombinant human bone morphogenetic protein-2 for the treatment of bone defects using non-viral scaffolds as gene delivery vehicles [65].

In view of their vast applications, robust scale-up platforms are needed for proper implementation of genetically engineered MSCs in clinical settings. As a proof-of-concept, we successfully transduced MSCs with lentiviral vectors expressing RFP. Transduction efficiencies obtained from our 3D model were promising, though 60% of ideal 2D monolayer controls, suggesting that microcapsules may restrict the contact between viral particles and cell membrane, preventing viral fusion and entry [66]. Importantly, transduction enhancing materials made a significant different in engineering MSCs in capsules. Different transduction enhancers have previously been investigated in the context of genetic manipulation of MSCs [35, 67]. These reagents mainly neutralize the natural surface charge of cells, enhancing viral adsorption by the presence of polycations. As expected, the use of a viral entry enhancer reagent significantly improved viral transduction in our system, and its use will be adopted in future studies. Moreover, to achieve higher and more consistent transduction efficiencies, further investigations are necessary, including, but not limited to, viral type, gene size, multiplicity of infection (MOI), encapsulation density and viral exposure time. Further analysis of proliferation and differentiation capabilities of transduced cells would also be of interest. Taken together, this study demonstrates the foundation for scaled-up genetic modification of MSCs and opens new possibilities to the use of 3D vertical-wheel systems in biomanufacturing.

## Conclusions

To meet the clinical demand of emerging MSC-based therapeutics, there remains a need to develop novel systems that will ensure consistent manufacturing and translation of these therapies. In this study, VitroGel-MSC cell-laden microcapsules were maintained in dynamic, suspension culture within a vertical-wheel bioreactor system using a fed-batch approach, while preserving critical quality attributes such as immunophenotype and multipotency after expansion and cell recovery. We have characterized critical parameters of the electrospraying encapsulation process such as seeding density, correlation of microcapsule output with hydrogel volume, applied voltage to fabricate cell-laden microcapsules of uniform size, and analyzed specific metabolic flux to better understand factors affecting this platform performance. We believe this study provides the foundations for bioprocess engineering of MSCs but can contribute throughout cell and gene therapy development.

## Authorship

Conceptualization, BP, MT; Execution of Experiments, MT, PJ, RB; Formal Analysis, MT, PJ, RB; Manuscript Preparation, MT, PJ, RB; Funding Acquisition, BP

## Acknowledgements

This research was conducted with support by under Contract number T0067 from the Advanced Regenerative Medicine Institute, BioFAB USA and Grant Nos. R01EB012521 (BP) and R01EB02872 (BP) awarded by the National Institutes of Health.

## Ethical Statements

The authors declare no conflict of interest. No ethical approval required.

Single donor bone marrow was donated for purchase from Lonza (Walkersville, MD, USA). Lonza obtained permission for its use in research applications by written informed consent.

## References

[1] A. Burr and B. Parekkadan (2019) Kinetics of MSC-based enzyme therapy for immunoregulation. Journal of Translational Medicine. 17: 263.

[2] M. Li, D. Khong, L. Y. Chin, A. Singleton and B. Parekkadan (2018) Therapeutic Delivery Specifications Identified Through Compartmental Analysis of a Mesenchymal Stromal Cell-Immune Reaction. Sci Rep. 8: 6816.

[3] Y. Zhang, M. Ravikumar, L. Ling, V. Nurcombe and S. M. Cool (2021) Age-Related Changes in the Inflammatory Status of Human Mesenchymal Stem Cells: Implications for Cell Therapy. Stem Cell Reports. 16: 694–707.

[4] M. Angelopoulou, E. Novelli, J. E. Grove, H. M. Rinder, C. Civin, L. Cheng and D. S. Krause (2003) Cotransplantation of human mesenchymal stem cells enhances human myelopoiesis and megakaryocytopoiesis in NOD/SCID mice. Exp Hematol. 31: 413–20.

[5] L. M. Ball, M. E. Bernardo, H. Roelofs, A. Lankester, A. Cometa, R. M. Egeler, F. Locatelli and W. E. Fibbe (2007) Cotransplantation of ex vivo expanded mesenchymal stem cells accelerates lymphocyte recovery and may reduce the risk of graft failure in haploidentical hematopoietic stem-cell transplantation. Blood. 110: 2764–7.

[6] J. F. Wang, Y. F. Wu, J. Harrintong and I. K. McNiece (2004) Ex vivo expansions and transplantations of mouse bone marrow-derived hematopoietic stem/progenitor cells. J Zhejiang Univ Sci. 5: 157–63.

[7] B. l. Lin, J. f. Chen, W. h. Qiu, K. w. Wang, D. y. Xie, X. y. Chen, Q. l. Liu, L. Peng, J. g. Li, Y. y. Mei, W. z. Weng, Y. w. Peng, H. j. Cao, J. q. Xie, S. b. Xie, A. P. Xiang and Z. l. Gao (2017) Allogeneic bone marrow– derived mesenchymal stromal cells for hepatitis B virus–related acute-on-chronic liver failure: A randomized controlled trial. Hepatology (Baltimore, Md.). 66: 209–219.

[8] P. Petrou, I. Kassis, N. Levin, F. Paul, Y. Backner, T. Benoliel, F. C. Oertel, M. Scheel, M. Hallimi, N. Yaghmour, T. B. Hur, A. Ginzberg, Y. Levy, O. Abramsky and D. Karussis (2020) Beneficial effects of autologous mesenchymal stem cell transplantation in active progressive multiple sclerosis. Brain (London, England : 1878). 143: 3574–3588.

[9] J. Kurtzberg, S. Prockop, P. Teira, H. Bittencourt, V. Lewis, K. W. Chan, B. Horn, L. Yu, J.-A. Talano, E. Nemecek, C. R. Mills and S. Chaudhury (2014) Allogeneic Human Mesenchymal Stem Cell Therapy (Remestemcel-L, Prochymal) as a Rescue Agent for Severe Refractory Acute Graft-versus-Host Disease in Pediatric Patients. Biology of blood and marrow transplantation. 20: 229–235.

[10] K. Zhao, R. Lou, F. Huang, Y. Peng, Z. Jiang, K. Huang, X. Wu, Y. Zhang, Z. Fan, H. Zhou, C. Liu, Y. Xiao, J. Sun, Y. Li, P. Xiang and Q. Liu (2015) Immunomodulation Effects of Mesenchymal Stromal Cells on Acute Graft-versus-Host Disease after Hematopoietic Stem Cell Transplantation. Biology of blood and marrow transplantation. 21: 97–104.

[11] K.-C. Moon, H.-S. Suh, K.-B. Kim, S.-K. Han, K.-W. Young, J.-W. Lee and M.-H. Kim (2019) Potential of Allogeneic Adipose-Derived Stem Cell-Hydrogel Complex for Treating Diabetic Foot Ulcers. Diabetes (New York, N.Y.). 68: 837–846.

[12] L. R. Braid, W. G. Hu, J. E. Davies and L. P. Nagata (2016) Engineered Mesenchymal Cells Improve Passive Immune Protection Against Lethal Venezuelan Equine Encephalitis Virus Exposure. Stem Cells Transl Med. 5: 1026–35.

[13] A. Allen, N. Vaninov, M. Li, S. Nguyen, M. Singh, P. Igo, A. W. Tilles, B. O’Rourke, B. L. K. Miller, B. Parekkadan and R. N. Barcia (2020) Mesenchymal Stromal Cell Bioreactor for Ex Vivo Reprogramming of Human Immune Cells. Scientific Reports. 10: 10142.

[14] J. A. Kink, M. H. Forsberg, S. Reshetylo, S. Besharat, C. J. Childs, J. D. Pederson, A. Gendron-Fitzpatrick, M. Graham, P. D. Bates, E. G. Schmuck, A. Raval, P. Hematti and C. M. Capitini (2019) Macrophages Educated with Exosomes from Primed Mesenchymal Stem Cells Treat Acute Radiation Syndrome by Promoting Hematopoietic Recovery. Biol Blood Marrow Transplant. 25: 2124–2133.

[15] M. Shao, Q. Xu, Z. Wu, Y. Chen, Y. Shu, X. Cao, M. Chen, B. Zhang, Y. Zhou, R. Yao, Y. Shi and H. Bu (2020) Exosomes derived from human umbilical cord mesenchymal stem cells ameliorate IL-6-induced acute liver injury through miR-455-3p. Stem Cell Res Ther. 11: 37.

[16] J. M. Hare, J. H. Traverse, T. D. Henry, N. Dib, R. K. Strumpf, S. P. Schulman, G. Gerstenblith, A. N. DeMaria, A. E. Denktas, R. S. Gammon, J. B. Hermiller, M. A. Reisman, G. L. Schaer and W. Sherman (2009) A Randomized, Double-Blind, Placebo-Controlled, Dose-Escalation Study of Intravenous Adult Human Mesenchymal Stem Cells (Prochymal) After Acute Myocardial Infarction. Journal of the American College of Cardiology. 54: 2277–2286.

[17] T. R. Olsen, K. S. Ng, L. T. Lock, T. Ahsan and J. A. Rowley (2018) Peak MSC-Are we there yet? Frontiers in medicine. 5: 178–178.

[18] J. Rowley, Abraham, E., Campbell, A., Brandwein, H., & Oh, S. (2012) Meeting Lot-Size Challenges of Manufacturing Adherent Cells for Therapy. BioProcess International. 10: 16–22.

[19] A. C. Schnitzler, A. Verma, D. E. Kehoe, D. Jing, J. R. Murrell, K. A. Der, M. Aysola, P. J. Rapiejko, S. Punreddy and M. S. Rook (2016) Bioprocessing of human mesenchymal stem/stromal cells for therapeutic use: Current technologies and challenges. Biochemical engineering journal. 108: 3–13.

[20] M. Teryek, A. Doshi, L. S. Sherman, P. Rameshwar, S. Jung and B. Parekkadan (2022) Clinical Manufacturing of Human Mesenchymal Stromal Cells using a Potency-Driven Paradigm. Current Stem Cell Reports. 8: 61–71.

[21] Q. A. Rafiq, K. Coopman, A. W. Nienow and C. J. Hewitt (2016) Systematic microcarrier screening and agitated culture conditions improves human mesenchymal stem cell yield in bioreactors. Biotechnol J. 11: 473–86.

[22] M. S. Croughan, D. Giroux, D. Fang and B. Lee (2016) Chapter 5 - Novel Single-Use Bioreactors for Scale-Up of Anchorage-Dependent Cell Manufacturing for Cell Therapies. pp. 105-139In: J. M. S. Cabral, C. Lobato de Silva, L. G. Chase and M. Margarida Diogo (eds.). Stem Cell Manufacturing. Elsevier, City.

[23] P. Silva Couto, M. C. Rotondi, A. Bersenev, C. J. Hewitt, A. W. Nienow, F. Verter and Q. A. Rafiq (2020) Expansion of human mesenchymal stem/stromal cells (hMSCs) in bioreactors using microcarriers: lessons learnt and what the future holds. Biotechnol Adv. 45: 107636.

[24] H. H. Tønnesen and J. Karlsen (2002) Alginate in Drug Delivery Systems. Drug Development and Industrial Pharmacy. 28: 621–630.

[25] M. S. Davis, I. Marrero-Berrios, I. Perez, C. P. Rabolli, P. Radhakrishnan, D. Manchikalapati, J. Schianodicola, H. Kamath, R. S. Schloss and J. Yarmush (2019) Alginate encapsulation for bupivacaine delivery and mesenchymal stromal cell immunomodulatory cotherapy. J Inflamm Res. 12: 87–97.

[26] S. Kumar, M. Kabat, S. Basak, J. Babiarz, F. Berthiaume and M. Grumet (2022) Anti-Inflammatory Effects of Encapsulated Human Mesenchymal Stromal/Stem Cells and a Method to Scale-Up Cell Encapsulation. In: Editor (ed.)^(eds.). Book Anti-Inflammatory Effects of Encapsulated Human Mesenchymal Stromal/Stem Cells and a Method to Scale-Up Cell Encapsulation, City.

[27] E. C. Stucky, J. Erndt-Marino, R. S. Schloss, M. L. Yarmush and D. I. Shreiber (2017) Prostaglandin E(2) Produced by Alginate-Encapsulated Mesenchymal Stromal Cells Modulates the Astrocyte Inflammatory Response. Nano Life. 7.

[28] C. L. Franco, J. Price and J. L. West (2011) Development and optimization of a dual-photoinitiator, emulsion-based technique for rapid generation of cell-laden hydrogel microspheres. Acta Biomater. 7: 3267–76.

[29] J. L. Wilson and T. C. McDevitt (2013) Stem cell microencapsulation for phenotypic control, bioprocessing, and transplantation. Biotechnol Bioeng. 110: 667–82.

[30] L. Huang, A. M. E. Abdalla, L. Xiao and G. Yang (2020) Biopolymer-Based Microcarriers for Three-Dimensional Cell Culture and Engineered Tissue Formation. International Journal of Molecular Sciences. 21: 1895.

[31] Z. Wang, D. Wu, J. Zou, Q. Zhou, W. Liu, W. Zhang, G. Zhou, X. Wang, G. Pei, Y. Cao and Z.-Y. Zhang (2017) Development of demineralized bone matrix-based implantable and biomimetic microcarrier for stem cell expansion and single-step tissue-engineered bone graft construction. Journal of Materials Chemistry B. 5: 62–73.

[32] C. Divieto and M. P. Sassi (2015) A first approach to evaluate the cell dose in highly porous scaffolds by using a nondestructive metabolic method. Future Sci OA. 1: Fso58.

[33] R. Bhatt, D. Ravi, A. M. Evens and B. Parekkadan (2022) Scaffold-mediated switching of lymphoma metabolism in culture. Cancer & Metabolism. 10: 15.

[34] E. Gabusi, E. Lenzi, C. Manferdini, P. Dolzani, M. Columbaro, Y. Saleh and G. Lisignoli (2022) Autophagy Is a Crucial Path in Chondrogenesis of Adipose-Derived Mesenchymal Stromal Cells Laden in Hydrogel. In: Editor (ed.)^(eds.). Book Autophagy Is a Crucial Path in Chondrogenesis of Adipose-Derived Mesenchymal Stromal Cells Laden in Hydrogel, City.

[35] K. Collon, M. C. Gallo, J. A. Bell, S. W. Chang, J. C. S. Rodman, O. Sugiyama, D. B. Kohn and J. R. Lieberman (2022) Improving Lentiviral Transduction of Human Adipose-Derived Mesenchymal Stem Cells. Hum Gene Ther. 33: 1260–1268.

[36] V. Neshati, S. Mollazadeh, B. S. Fazly Bazzaz, A. A. de Vries, M. Mojarrad, H. Naderi-Meshkin, Z. Neshati and M. A. Kerachian (2018) Cardiomyogenic differentiation of human adipose-derived mesenchymal stem cells transduced with Tbx20-encoding lentiviral vectors. J Cell Biochem. 119: 6146–6153.

[37] A. A. Mangi, N. Noiseux, D. Kong, H. He, M. Rezvani, J. S. Ingwall and V. J. Dzau (2003) Mesenchymal stem cells modified with Akt prevent remodeling and restore performance of infarcted hearts. Nat Med. 9: 1195–201.

[38] S. Kumar and S. Ponnazhagan (2007) Bone homing of mesenchymal stem cells by ectopic alpha 4 integrin expression. Faseb j. 21: 3917–27.

[39] A. Amari, M. Ebtekar, S. M. Moazzeni, M. Soleimani, L. Mohammadi Amirabad, M. T. Tahoori and M. Massumi (2015) In Vitro Generation of IL-35-expressing Human Wharton’s Jelly-derived Mesenchymal Stem Cells Using Lentiviral Vector. Iran J Allergy Asthma Immunol. 14: 416–26.

[40] Y. Zhu, M. Cheng, Z. Yang, C. Y. Zeng, J. Chen, Y. Xie, S. W. Luo, K. H. Zhang, S. F. Zhou and N. H. Lu (2014) Mesenchymal stem cell-based NK4 gene therapy in nude mice bearing gastric cancer xenografts. Drug Des Devel Ther. 8: 2449–62.

[41] T. K. Goh, Z. Y. Zhang, A. K. Chen, S. Reuveny, M. Choolani, J. K. Chan and S. K. Oh (2013) Microcarrier culture for efficient expansion and osteogenic differentiation of human fetal mesenchymal stem cells. Biores Open Access. 2: 84–97.

[42] T. R. Heathman, A. Stolzing, C. Fabian, Q. A. Rafiq, K. Coopman, A. W. Nienow, B. Kara and C. J. Hewitt (2016) Scalability and process transfer of mesenchymal stromal cell production from monolayer to microcarrier culture using human platelet lysate. Cytotherapy. 18: 523–35.

[43] M. Hervy, J. L. Weber, M. Pecheul, P. Dolley-Sonneville, D. Henry, Y. Zhou and Z. Melkoumian (2014) Long term expansion of bone marrow-derived hMSCs on novel synthetic microcarriers in xeno-free, defined conditions. PloS one. 9: e92120–e92120.

[44] E. Jain, K. M. Scott, S. P. Zustiak and S. A. Sell (2015) Fabrication of Polyethylene Glycol-Based Hydrogel Microspheres Through Electrospraying: Fabrication of Polyethylene Glycol-Based Hydrogel. Macromolecular materials and engineering. 300: 823–835.

[45] J. Mumaw, E. T. Jordan, C. Sonnet, R. M. Olabisi, E. A. Olmsted-Davis, A. R. Davis, J. F. Peroni, J. L. West, F. West, Y. Lu and S. L. Stice (2012) Rapid Heterotrophic Ossification with Cryopreserved Poly(ethylene glycol-) Microencapsulated BMP2-Expressing MSCs. Int J Biomater. 2012: 861794.

[46] D. Perera, M. Medini, D. Seethamraju, R. Falkowski, K. White and R. M. Olabisi (2018) The effect of polymer molecular weight and cell seeding density on viability of cells entrapped within PEGDA hydrogel microspheres. J Microencapsul. 35: 475–481.

[47] T. R. J. Heathman, Q. A. Rafiq, A. K. C. Chan, K. Coopman, A. W. Nienow, B. Kara and C. J. Hewitt (2016) Characterization of human mesenchymal stem cells from multiple donors and the implications for large scale bioprocess development. Biochemical engineering journal. 108: 14–23.

[48] G. Siegel, T. Kluba, U. Hermanutz-Klein, K. Bieback, H. Northoff and R. Schäfer (2013) Phenotype, donor age and gender affect function of human bone marrow-derived mesenchymal stromal cells. BMC medicine. 11: 146–146.

[49] O. Gryshkov, D. Pogozhykh, H. Zernetsch, N. Hofmann, T. Mueller and B. Glasmacher (2014) Process engineering of high voltage alginate encapsulation of mesenchymal stem cells. Materials Science and Engineering: C. 36: 77–83.

[50] A. S. Qayyum, E. Jain, G. Kolar, Y. Kim, S. A. Sell and S. P. Zustiak (2017) Design of electrohydrodynamic sprayed polyethylene glycol hydrogel microspheres for cell encapsulation. Biofabrication. 9: 025019.

[51] J. Lembong, R. Kirian, J. D. Takacs, T. R. Olsen, L. T. Lock, J. A. Rowley and T. Ahsan (2020) Bioreactor Parameters for Microcarrier-Based Human MSC Expansion under Xeno-Free Conditions in a Vertical-Wheel System. Bioengineering (Basel). 7: 73.

[52] Q. A. Rafiq, S. Ruck, M. P. Hanga, T. R. J. Heathman, K. Coopman, A. W. Nienow, D. J. Williams and C. J. Hewitt (2018) Qualitative and quantitative demonstration of bead-to-bead transfer with bone marrow-derived human mesenchymal stem cells on microcarriers: Utilising the phenomenon to improve culture performance. Biochemical Engineering Journal. 135: 11–21.

[53] A. T.-L. Lam, J. Li, A. K.-L. Chen, S. Reuveny, S. K.-W. Oh and W. R. Birch (2014) Cationic Surface Charge Combined with Either Vitronectin or Laminin Dictates the Evolution of Human Embryonic Stem Cells/Microcarrier Aggregates and Cell Growth in Agitated Cultures. Stem Cells and Development. 23: 1688–1703.

[54] M. Al-Rubeai, R. P. Singh, M. H. Goldman and A. N. Emery (1995) Death mechanisms of animal cells in conditions of intensive agitation. Biotechnology and Bioengineering. 45: 463–472.

[55] C. Selden, J. Bundy, E. Erro, E. Puschmann, M. Miller, D. Kahn, H. Hodgson, B. Fuller, J. Gonzalez-Molina, A. Le Lay, S. Gibbons, S. Chalmers, S. Modi, A. Thomas, P. Kilbride, A. Isaacs, R. Ginsburg, H. Ilsley, D. Thomson, G. Chinnery, N. Mankahla, L. Loo and C. W. Spearman (2017) A clinical-scale BioArtificial Liver, developed for GMP, improved clinical parameters of liver function in porcine liver failure. Sci Rep. 7: 14518.

[56] S. Swioklo, P. Ding, A. W. Pacek and C. J. Connon (2017) Process parameters for the high-scale production of alginate-encapsulated stem cells for storage and distribution throughout the cell therapy supply chain. Process Biochemistry. 59: 289–296.

[57] D. Schop, F. W. Janssen, L. D. van Rijn, H. Fernandes, R. M. Bloem, J. D. de Bruijn and R. van Dijkhuizen-Radersma (2009) Growth, metabolism, and growth inhibitors of mesenchymal stem cells. Tissue Eng Part A. 15: 1877–86.

[58] V. H. Pattasseril J, Lock L, Rowley JA (2013) Downstream Technology Landscape for Large-Scale Therapeutic Cell Processing. BioProcess International. 11: 38–47.

[59] G. M. Pigeau, E. Csaszar and A. Dulgar-Tulloch (2018) Commercial Scale Manufacturing of Allogeneic Cell Therapy. Frontiers in Medicine. 5.

[60] J. F. Welter, L. A. Solchaga and K. J. Penick (2007) Simplification of aggregate culture of human mesenchymal stem cells as a chondrogenic screening assay. Biotechniques. 42: 732, 734-7.

[61] D. M. Panchision, H.-L. Chen, F. Pistollato, D. Papini, H.-T. Ni and T. S. Hawley (2007) Optimized Flow Cytometric Analysis of Central Nervous System Tissue Reveals Novel Functional Relationships Among Cells Expressing CD133, CD15, and CD24. Stem Cells. 25: 1560–1570.

[62] J. L. Santos, D. Pandita, J. Rodrigues, A. P. Pêgo, P. L. Granja and H. Tomás (2011) Non-viral gene delivery to mesenchymal stem cells: methods, strategies and application in bone tissue engineering and regeneration. Curr Gene Ther. 11: 46–57.

[63] F. Marofi, G. Vahedi, A. Biglari, A. Esmaeilzadeh and S. S. Athari (2017) Mesenchymal Stromal/Stem Cells: A New Era in the Cell-Based Targeted Gene Therapy of Cancer. Front Immunol. 8: 1770.

[64] P. Lin, Y. Lin, D. P. Lennon, D. Correa, M. Schluchter and A. I. Caplan (2012) Efficient lentiviral transduction of human mesenchymal stem cells that preserves proliferation and differentiation capabilities. Stem Cells Transl Med. 1: 886–97.

[65] S. Andrews, A. Cheng, H. Stevens, M. T. Logun, R. Webb, E. Jordan, B. Xia, L. Karumbaiah, R. E. Guldberg and S. Stice (2019) Chondroitin Sulfate Glycosaminoglycan Scaffolds for Cell and Recombinant Protein-Based Bone Regeneration. Stem Cells Transl Med. 8: 575–585.

[66] D. S. Dimitrov (2004) Virus entry: molecular mechanisms and biomedical applications. Nat Rev Microbiol. 2: 109–22.

[67] F. Amadeo, V. Hanson, P. Murray and A. Taylor (2022) DEAE-Dextran Enhances the Lentiviral Transduction of Primary Human Mesenchymal Stromal Cells from All Major Tissue Sources Without Affecting Their Proliferation and Phenotype. Mol Biotechnol.

