## Supplemental Figure 1 for "3D Microcapsules for Human Bone Marrow Derived Mesenchymal Stem Cell Biomanufacturing in a Vertical-Wheel Bioreactor"

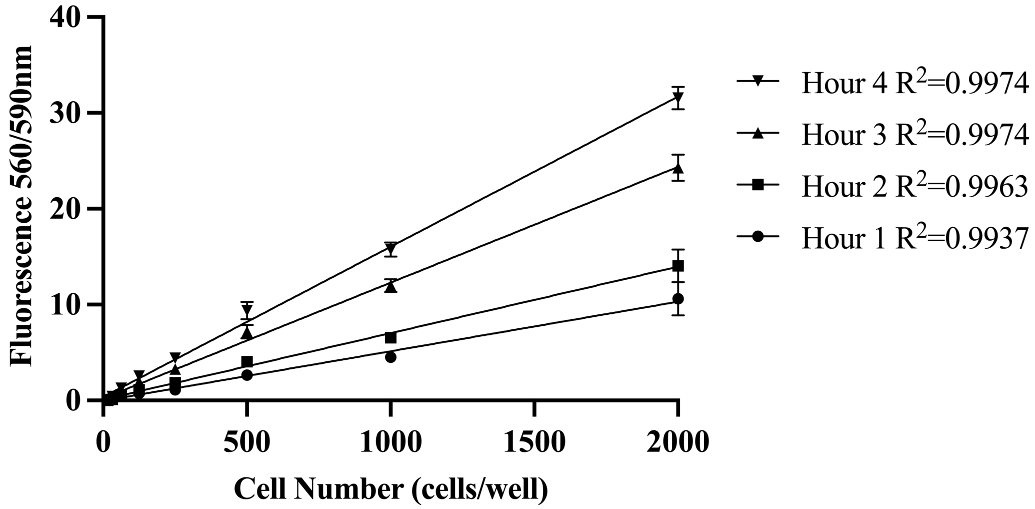


**Fig. S1.** Evaluation of CellTiter-Blue incubation time on fluorescent signal output. Serial two-fold dilution of MSCs prepared at 100 uL/well in a 96-well plate.
