## Supplementary figures and images for "3D Microcapsules for Human Bone Marrow Derived Mesenchymal Stem Cell Biomanufacturing in a Vertical-Wheel Bioreactor"

### Supplemental Figure 2

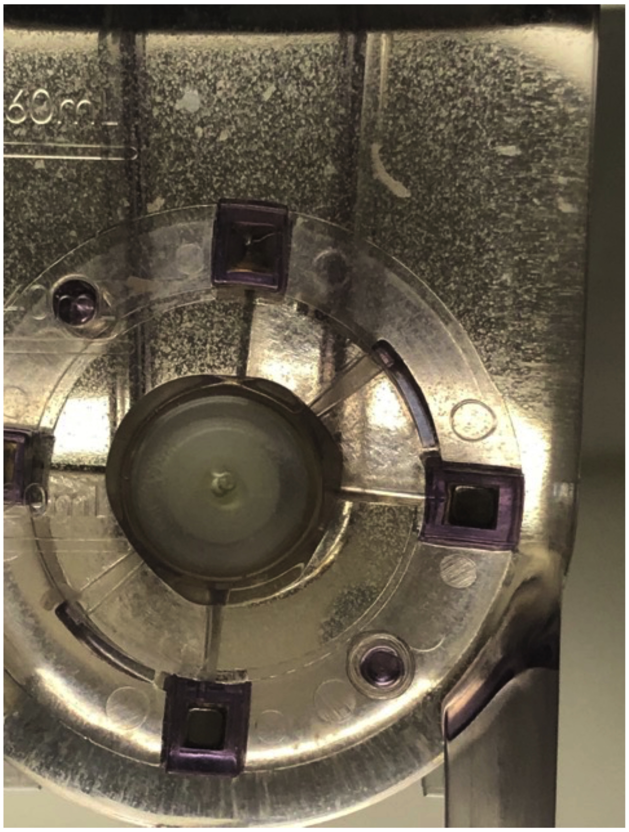

**Fig. S2.** Day 6 image of VitroGel-MSC microcapsules suspended within a vertical-wheel bioreactor.
