## Supplemental Figure 3 for "3D Microcapsules for Human Bone Marrow Derived Mesenchymal Stem Cell Biomanufacturing in a Vertical-Wheel Bioreactor"

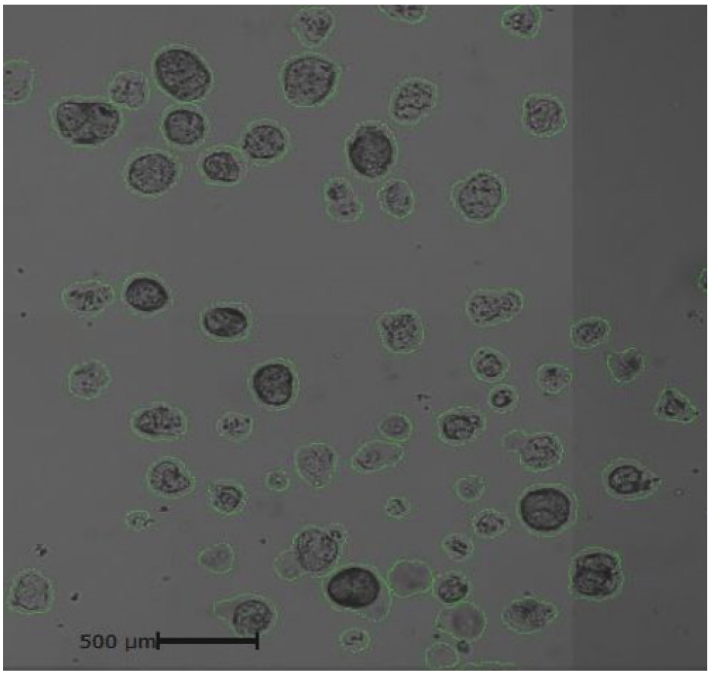


**Fig. S3.** Representative image of Celigo Image Cytometer settings used to characterize VitroGel-MSC microcapsule size distribution.
