## Supplemental Figure 4 for "3D Microcapsules for Human Bone Marrow Derived Mesenchymal Stem Cell Biomanufacturing in a Vertical-Wheel Bioreactor"

### VitroGel-MSC Microcapsules

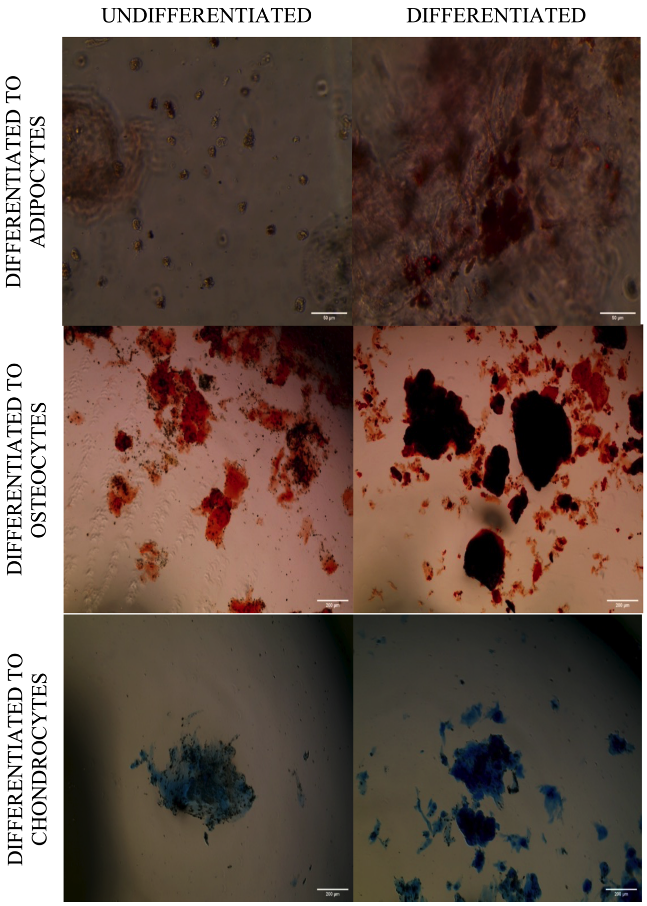


**Fig. S4.** Day 21 images of MSCs differentiated into adipocytes, osteocytes, and chondrocytes within VitroGel-MSC microcapsules.
