## Supplemental Figure 5 for "3D Microcapsules for Human Bone Marrow Derived Mesenchymal Stem Cell Biomanufacturing in a Vertical-Wheel Bioreactor"

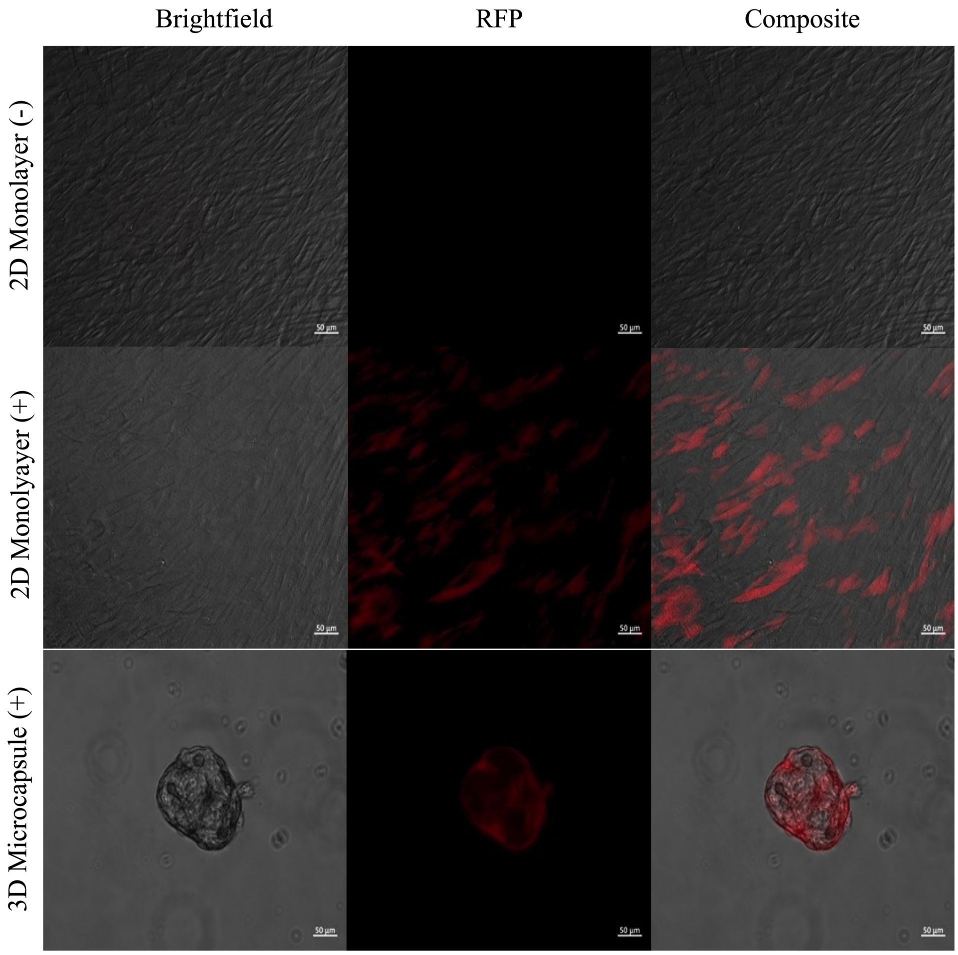


**Fig. S5.** Day 6 single channel and composite images evaluating MSC expression of RFP in 2D Monolayer controls and 3D VitroGel-MSC microcapsules. MSCs within VitroGel-MSC microcapsules were co-electrosprayed with lentiviral particles at MOI 50.
